# Peripheral sTREM2-related inflammatory activity alterations in early stage Alzheimer’s disease

**DOI:** 10.1101/2021.08.12.455916

**Authors:** Grace E. Weber, Maria Khrestian, Elizabeth D. Tuason, Yvonne Shao, Jagan Pillai, Stephen Rao, Hao Feng, Yadi Zhou, Feixiong Cheng, Tara M. DeSilva, Shaun Stauffer, James B. Leverenz, Lynn M. Bekris

**Author notes:** **Corresponding Author: Dr. Lynn M. Bekris**, (216)217-1791. Funding: National Institute of Aging (R01AG066707 and 3R01AG066707-01S1 (analysis and interpretation of data) AG063870 (collection, analysis and interpretation of data), P30 AG062428 (collection), R01 AG022304 (collection), Cleveland Clinic Center of Excellence Award (interpretation of data), Jane and Lee Seidman Fund (collection, analysis and interpretation of data).

## Abstract

Alzheimer’s disease (AD) has been linked to multiple immune system genetic variants, implicating potential broad alterations in inflammatory profiles in the disease. Triggering receptor expressed on myeloid cells 2 (TREM2) genetic variants are risk factors for AD and other neurodegenerative diseases. A soluble TREM2 isoform (sTREM2) is elevated in cerebrospinal fluid in the early stages of AD suggesting it may be a biomarker of progressive alterations in immune response to AD-related pathology. Multiple studies have reported an altered peripheral immune response in AD. However, less is known about the relationship between plasma sTREM2 and the altered peripheral immune response in AD. The objective of this exploratory study was to examine the relationship between sTREM2 and inflammatory activity in human participants defined by clinically characterized cognitive symptoms and groups defined by the cerebrospinal fluid biomarkers <u>a</u>myloid beta, phosphorylated <u>t</u>au, and <u>n</u>eurodegeneration (NIA-AA Research Framework: “ATN continuum”.) The hypothesis of this exploratory study was that sTREM2 related inflammatory activity differs by AD stage. We observed different patterns of inflammatory activity across disease groups and ATN categories that implicates peripheral sTREM2 related inflammatory activity as altered in the early stages of AD. Notably, fractalkine showed a significant relationship with sTREM2 across different analyses in the control groups that was lost as disease progressed, and fractalkine, IL-5 and IL-17A were decreased in AD. These preliminary data provide important support to the hypothesis that sTREM2-related inflammatory activity is a stage-specific biomarker of AD progression, providing the groundwork for future studies and therapeutic strategies.

## Introduction

Alzheimer’s disease is a devastating neurodegenerative disorder that impacts about 5.8 million Americans and is the 6^th^ leading cause of death in the United States (1). Typically, Alzheimer’s disease (AD) - related pathology, which includes amyloid beta (Aβ) plaques and tau neurofibrillary tangles, begin to accumulate years prior to the onset of dementia symptoms. This results in a delay in diagnosing AD until after neurodegeneration has progressed. In 2021 the FDA approved Aduhelm (aducanumab) for Alzheimer’s disease treatment, but there is currently no validated blood-based biomarker to predict AD risk or onset (2, 3). Clinically, AD is diagnosed using cerebrospinal fluid (CSF) measures of Aβ and tau, neurological testing and brain imaging when available. However, AD can only be definitively diagnosed and distinguished from other neurodegenerative disease and dementia types upon autopsy based on brain pathology (4, 5).

Stages of AD progression are primarily based on clinical signs and symptoms; a more recent classification method is based on CSF biomarkers. In the classic symptomatic classification, asymptomatic individuals are classified as “cognitively normal,” often in the absence of testing for CSF biomarkers and brain pathology. Cognitively normal (CN) individuals have no presentation of the symptoms of dementia such as memory loss, or do not present the symptoms in a way that is more severe than expected age-related forgetfulness. The next stage is mild cognitive impairment (MCI), in which cognition has declined to an extent greater than expected for one’s age but without having Alzheimer’s disease or other diagnoses. A form of MCI thought to progress into AD dementia is referred to as amnestic MCI (6). Finally, a person can progress to AD dementia, of which there are also progressively mild, moderate, and severe forms (7). The ATN continuum is a second method of defining stages of neurodegenerative diseases, including AD dementia, and is based on measures of CSF biomarker proteins. “ATN” refers to “A” amyloid β (Aβ), “T” phosphorylated tau (p-Tau), and “N” neurodegeneration. Aβ is normally 40kD (Aβ40), while a pathogenic form is 42kD (Aβ42). The ratio of pathogenic: normal Aβ, Aβ42/ Aβ40, is used as a criterion for diagnosis. Similarly, the tau protein can be phosphorylated (p-Tau), which is the form most commonly present in neurofibrillary tangles. The ATN continuum defines individuals as A-T-N- if they have no CSF Aβ or tau abnormalities, A+T-N- if they have only altered CSF Aβ42/ Aβ40 ratios, and progresses to A+T+N+, which is more typical in AD than CN individuals (8). Few studies to date have directly compared peripheral inflammatory markers within people diagnosed via both methods described.

AD is a disease of the brain, but there is evidence that peripheral inflammation contributes to brain inflammation and AD pathology. Several inflammatory disorders are risk factors for AD including obesity, metabolic syndrome, traumatic brain injury, and chronic periodontitis development (9). It has been suggested that recurrent inflammatory events over one’s life including infections, ischemia, and free radical exposure, parallel to the accumulation of Aβ, can increase the risk of AD development due to the repeated cycle of activated immune cells both in the peripheral and the central nervous system (10). The mechanism underlying the link between peripheral inflammation and AD is not fully understood.

In order to better understand the role of peripheral inflammation and AD, studies have measured levels of circulating inflammatory factors. Importantly, there is evidence that immune system dysregulation can also contribute to risk of AD, and it may be the lack of a proper immune system reaction that makes people susceptible to repeated infections and consequently AD. For example, a retrospective review of hospital data found that participants admitted to the hospital with autoimmune disorders were more likely to be diagnosed with AD, and 18 of the 25 autoimmune disorders that were part of the study showed significant correlations with dementia (11). Similarly, age-related decline in immune system function,\ referred to as immunosenescence, may contribute to risk of AD (12). Meta-analyses of inflammatory factors in people with AD compared to controls often show mixed results which may be a result of differing study groups and confounding factors including age and *APOE* e4 status (13–15).

One immune receptor associated with AD is triggering receptor expressed on myeloid cells 2 (TREM2). Normally, TREM2 functions as a pattern recognition receptor in the innate immune response. It is expressed on microglia in the brain and myeloid cells in peripheral blood. Several loss-of-function mutations in *TREM2* have been shown to increase AD risk (16, 17). The R47H *TREM2* mutation, results in decreased function of the receptor (16). Another mutation associated with increased AD risk is H157Y. It leads to less full-length cell-surface TREM2 and more of the cleaved version, soluble TREM2 (sTREM2), which is detectible in CSF and plasma (18, 19). Increased CSF sTREM2 in MCI and AD has been described in multiple studies while results from plasma are less clear (20, 21). It is still unclear whether sTREM2 in the CSF or plasma indicates increased expression of membrane-bound TREM2 or increased cleavage of sTREM2 or both.

Since there is a relationship between sTREM2 and AD and evidence of inflammatory factor alterations in AD, the aim of this exploratory investigation was to investigate the relationship between plasma sTREM2 and peripheral inflammatory activity in AD. The primary hypothesis is that alterations in plasma sTREM2 is related to altered peripheral inflammatory activity in AD, including the early stages of the disease, defined by mild cognitive impairment or early AD pathological categories (ATN category). This is important because it will define the poorly understood relationship between the broader inflammatory response and TREM2 in both the MCI stage and in AD. In addition, it will determine whether TREM2-related inflammatory activity is a feasible biomarker candidate for discerning AD subtypes or stages.

## Materials and Methods

### Participant recruitment and study design

Participants defined as cognitively normal (CN) controls (n=88), amnestic mild cognitive impairment (MCI; n=37) and Alzheimer’s disease dementia (AD; n=75) donated biospecimens under the Lou Ruvo Center for Brain Health Aging and Neurodegenerative Disease Biobank (LRCBH-Biobank) and the Cleveland Alzheimer’s Disease Research Center (CADRC) protocols approved by the Cleveland Clinic Institutional Review Board. Participants were all over 55 years of age. Venipunctures were performed after overnight fasting for the collection of whole blood. Plasma was isolated from lavender-top EDTA tubes and stored at −80°C. CSF was aliqouted into amber tubes and immediately frozen and stored at −80°C as previously described (22).

### Neurological testing, diagnosis, and consensus

All study participants underwent cognitive testing upon enrollment in Cleveland Clinic LRCBH-biobank or CADRC. The participants were reviewed by neurologists in a formal consensus panel to assign diagnostic groups. Only the groups CN (n=88), MCI without AD or other diagnoses (n=37) and AD dementia (n=75) were included in this study (Table I). A subset of this group had both a blood draw and a lumbar puncture for collection of CSF for which AD-related biomarkers Aβ40, Aβ42, total tau (t-Tau), and phosphorylated-tau 181 (p-Tau) were measured for research studies (CN: n=27, MCI: n= 27, AD: n= 65) (Table I).

**Table I:** Cohort demographics, clinical chemistry, and AD-related biomarkers. The cohort was comprised of cognitively normal (CN), mild cognitive impairment (MCI) and Alzheimer’s disease (AD) participants, ages 55-86 years. Age ranges and average ages per group are shown, and did not differ between groups by Kruskal-Wallis test. Proportions of males participants were similar between groups but *APOE* 4+ alleles were more prevalent in MCI And AD than CN (by Fisher’s exact test). AD showed significantly higher levels of WBC compared to MCI and lower basophil percentages compared to CN, while CN had higher erythrocyte sedimentation rates compared to MCI (by Kruskal-Wallis). Finally, AD-related biomarkers were significantly different between disease groups as expected. Significant differences are noted in bold.

|  | CN | MCI | AD | CN vs<br>MCI | CN vs<br>AD | MCI vs<br>AD |
| --- | --- | --- | --- | --- | --- | --- |
| **Cohort Description** |  |  |  |  |  |  |
| n = | 88 | 37 | 75 |  |  |  |
| Age range (years) | 55-84 | 55-83 | 56-86 |  |  |  |
| Average age (years) | 66.5 | 68.6 | 66.4 | 0.3576 | >0.9999 | 0.2091 |
| % Male | 44.3 | 59.5 | 49.3 | 0.1698 | 0.5332 | 0.3241 |
| % *APOE*4+ | 38.6 | 67.6 | 66.7 | **0.0035** | **0.0005** | >0.9999 |
| **Clinical chemistry** |  |  |  |  |  |  |
| Complete Blood Counts: n = | 49 | 33 | 60 |  |  |  |
| White Blood Cells mean (SD) | 5.5 (1.4) | 4.9 (1.1) | 6.0 (2.1) | 0.2234 | >0.9999 | **0.0239** |
| Neutrophil % mean (SD) | 60.7 (8.4) | 61.0 (9.1) | 62.5 (10.8) | >0.9999 | 0.5727 | >0.9999 |
| Lymphocyte % mean (SD) | 27.7 (6.8) | 26.9 (9.1) | 26.3 (8.9) | >0.9999 | 0.6040 | >0.9999 |
| Monocyte % mean (SD) | 8.3 (1.8) | 8.9 (1.8) | 8.5 (3.2) | 0.1540 | >0.9999 | 0.1069 |
| Basophil % mean (SD) | 0.53 (0.28) | 0.45 (0.23) | 0.39 (0.24) | 0.8922 | **0.0309** | 0.6943 |
| Erythrocyte Sedimentation Rate: n = | 48 | 33 | 59 |  |  |  |
| Erythrocyte Sedimentation Rate Mean (SD) | 12.2 (9.1) | 7.5 (6.1) | 8.7 (6.0) | **0.0188** | 0.4160 | 0.3817 |
| C-reactive Protein: n = | 46 | 32 | 53 |  |  |  |
| C-reactive Protein Mean (SD) | 1.9 (2.0) | 3.8 (10.9) | 2.5 (4.7) | >0.9999 | >0.9999 | 0.7810 |
| Blood Brain Barrier Integrity: n = | 13 | 22 | 35 |  |  |  |
| Blood Brain Barrier Integrity Mean (SD) | 0.006 | 0.008 | 0.006 | 0.5887 | >0.9999 | 0.1321 |
| **AD-related Biomarkers (ATN)** |  |  |  |  |  |  |
| n = | 27 | 27 | 65 |  |  |  |
| CSF AB 42/40 (A) mean (SD) | 0.17 (0.05) | 0.12 (0.09) | 0.09 (0.03) | **0.0027** | **<0.0001** | **0.0019** |
| CSF pTau-181 (pg/ml) (T) mean (SD) | 44.8 (21.4) | 98.3 (75.5) | 150.8 (111.8) | **0.0232** | **<0.0001** | **0.0021** |
| CSF tTau (pg/ml) (N) mean (SD) | 379.9 (225.2) | 576.8 (431.1) | 763.0 (458.1) | >0.9999 | **0.0004** | **0.0098** |

### Clinical testing

Complete blood counts (CBC) were performed on n=49 CN, n=33 MCI, and n=60 AD participants via microscopy by trained Cleveland Clinic technicians blinded to sample group. Lumbar punctures were performed for the collection of CSF from a subset of participants (CN: n=13, MCI: n= 22, AD: n= 35) to assess blood brain barrier integrity (Table I). Blood and CSF samples were drawn on the same date in each participant to allow for direct comparison of clinical markers and study markers.

### APOE genotyping

Genotyping of apolipoprotein E (*APOE*) was performed from blood samples using the 7500 Real Time PCR System and TaqMan SNP Genotyping Assays (rs429358, rs7412) (Thermo Fisher Scientific) as previously described (23).

### Plasma sTREM2

Plasma soluble TREM2 (sTREM2) levels were measured using a Luminex 200 3.1 xPONENT System (EMD Millipore, Chicago, IL, USA) and a custom detection method designed to capture the soluble portion of TREM2 protein as previously described (24). Briefly, a capture antibody bound to MagPlex beads binds sTREM2 (R&D #MAB1828 human TREM2 antibody monoclonal mouse IgG_2B_ Clone #263602; Immunogen His19-Ser174). A biotinylated antibody with a SAPE conjugate was used for detection (R&D: #BAF1828; human TREM2 biotinylated antibody; antigen affinity-purified polyclonal goat IgG; Immunogen His19-Ser174).

### Plasma inflammatory markers

A panel of 38 plasma inflammatory factors (cytokines, chemokines, and growth factors) were measured with a human cytokine/chemokine panel utilizing Luminex 200 xMap technology and the MILLIPLEX MAP ^®^ multiplex kit (Luminex xMAP technology; EMD Millipore, Chicago, IL, USA, kit HCYTMAG60PMX41BK) following the manufacturer’s instructions for analyte detection in human plasma. This panel was selected because it included both previously-studied cytokines and cytokines without previous study with sTREM2 in AD. The inflammatory markers in the panel were: Epidermal Growth Factor (EGF), Fibroblast Growth Factor 2 (FGF-2), eotaxin, Transforming Growth Factor alpha (TGF-α), Granulocyte-colony stimulating factor (G-CSF), FMS-like tyrosine kinase 3 ligand (Flt-3L), Granulocyte-Macrophage Colony Stimulating Factor (GM-CSF), Fractalkine (also known as CX3CL1), interferon (IFN) α2 (IFNα2), IFNγ, growth-regulated oncogene (GRO), interleukin (IL) 10, (IL-10), Monocyte chemotactic protein-3 (MCP-3, also known as CCL7), IL-12 40kDa (IL-12p40), Macrophage-derived chemokine (MDC), IL-12 70kDa (IL-12P70), IL-13, IL-15, soluble CD40-ligand (sCD40L), IL-17A, IL-1 receptor agonist (IL-1RA), IL-1α, IL-9, IL-1β, IL-2, IL-3, IL-4, IL-5, IL-6, IL-7, IL-8, interferon-gamma inducible protein 10kDa (IP-10), MCP-1 (also known as CCL2), macrophage inflammatory protein (MIP) 1α (MIP-1α also known as CCL3), MIP-1β (also known as CCL4), tumor necrosis factor (TNF) alpha (TNFα), TNFβ, and Vascular Endothelial Growth Factor (VEGF).

### Quality control

Plasma sTREM2 and inflammatory markers were measured in plasma on plates that also contained buffer-alone background wells and serial dilutions of analytes in order to generate a standard curve. Plates measuring inflammatory markers also contained quality control proteins. Each plate was evaluated to ensure that the standards and quality controls ran in the expected ranges. Samples from different disease groups were run across each plate to control for batch effects. Mean fluorescence intensities (MFI) of analytes were calculated and averaged between duplicate samples. Our analyses compared MFI rather than calculated concentration because many participants had low levels of cytokines, which were undetectable by the standard curve interpolation. Utilization of MFI values allowed inclusion of all participants, including the samples at the low range of detection, to avoid biasing the results towards participants with higher values (25–27).

### CSF biomarkers

CSF Aβ40, Aβ42, total tau (t-Tau), and phosphorylated-tau 181 (p-Tau) were measured according to manufacturer specifications (Luminex xMAP technology; EMD Millipore, Chicago, IL, USA: HNABTMAG-68K), modified by a 1:10 dilution of CSF. Each Aβ and Tau kit comes with an Aβ and Tau standard, as well as Aβ and Tau quality controls. The kit provides the expected concentrations of each working standard as well as each of the quality controls. The standards, controls, and CSF samples are all run in duplicate. If the coefficient of variation for any of the replicate wells is greater than 25% or if both replicate wells have a bead count of less than 35 beads for a given analyte, the results are repeated or excluded for that analyte.

### ATN groups

CSF measurements of Aβ40, Aβ42, t-Tau, and p-Tau were performed as described above. Participants were categorized into ATN groups based on the following criteria: A+ indicates participants were in the lower 10^th^ percentile of all Aβ42/Aβ40 ratios within CN group in the cohort, T+ indicates participants were in the upper 90^th^ percentile of all phosphorylated tau protein concentrations within the CN group in the cohort, and N+ indicates participants were in the upper 90^th^ percentile of total tau protein concentrations within CN group in the cohort. Cutoff values generated by percentile analysis were as follows: A+ ≤ Aβ42/Aβ40 0.139; T+ p-Tau≥ 65.5pg/ml; N+ t-Tau ≥ 695.9pg/ml. These cutoff values were tested in contingency tables with numbers of CN and AD participants. The cutoffs were selected because they had the highest sensitivity and specificity in our cohort, compared to cutoffs set at 20^th^/80^th^ percentiles, 15^th^/85^th^ percentiles, and 5^th^/95^th^ percentiles (for Aβ42/Aβ40, and p-Tau and T-Tau, respectively) as well as concentrations previously published (28–30). Using the 10^th^/90^th^ percentiles and cutoffs listed above our cutoffs had Aβ42/Aβ40: sensitivity=0.963, specificity=0.901 (p<0.001 by Fisher’s exact test); p-Tau: sensitivity=0.722, specificity=0.887 (p<0.001 by Fisher’s exact test); and t-Tau: sensitivity=0.426, specificity=0.811 (p=0.0266 by Fisher’s exact test).

### Plasma sTREM2 Tertiles

We characterized participants into sTREM2 tertiles in order to compare groups with the highest versus lowest levels of plasma sTREM2. Levels of sTREM2 from participants were sorted from lowest to highest, regardless of disease status and ATN group, and the results were divided into thirds. Tertile 1 (n=67) refers to the lowest sTREM2 levels, and Tertile 3 (n=67) refers to the highest sTREM2 levels. Tertile 2 (n=66) was not included in the tertile analyses. Tertiles were used rather than quartiles or quintiles in order to maximize the sample size while still analyzing two distinct groups of sTREM2 levels.

### Statistical Analyses

CN, MCI, and AD demographics, plasma inflammatory marker, and plasma sTREM2 were compared between groups using Kruskal-Wallis tests with Dunn’s test for multiple comparisons, after outlier removal using the ROUT method (Q=1%). ATN groups were compared by Mann-Whitney tests along the ATN continuum (A-T-N- vs A+T-N-, A+T-N- vs A+T+N-, and A+T+N- vs A+T+N), after outlier removal using the ROUT method (Q=1%). Nonparametric tests were used because the data did not pass normality tests using the Anderson-Darling and D’Agostino-Pearson methods. Sensitivity and specificity of ATN group cutoffs were analyzed by Fisher’s exact tests and contingency tables of sTREM2 tertiles in disease groups were analyzed by Chi square tests. These data were analyzed, and violin plots and bar graphs were generated using GraphPad Prism version 9.0.2 (GraphPad Software, San Diego, CA). Using R (R version 3.6.1), the “lm” function was used to create a single linear regression model for each cytokine. The standardized beta coefficient was then obtained by applying the “standardize_parameters” function from the “*effectsize*” package to these models (31). The “cor” function was used to determine Spearman correlations. Correlograms were made using the resulting spearman correlation matrix with the “corrplot” function in the “*corrplot*” package (32) and arranged according to hierarchical clustering using the Ward’s method. The p-values were added using results from the “rcorr” function from the “Hmisc” function as a parameter in the corrplot function (33). To compare the graphs of correlation matrix, the Steiger test was applied to the correlation matrices with the “cortest.normal” function from the psych package (34). A power analysis utilizing analyte data from a small pilot study indicated a minimum sample size of n=25 per group was necessary to achieve a significance of 0.05 and a power of 0.8 for these analyses.

### Co-expression network visualization

A co-expression network was generated for each group based on the Spearman’s rank correlation coefficient ρ. For inflammatory factors, Spearman r > 0.8 and p < 0.05 was used as cutoff for co-expressions; for sTREM2 vs. other inflammatory factors, Spearman r > 0 and p < 0.05 was used as a cutoff. The co-expression networks were visualized by Gephi 0.9.2 (35).

## Results

### Cohort demographics and clinical measurements

Characteristics of the study populations were compared between cognitively normal (CN) controls (n=88), individuals with MCI (n=37), and participants with AD (n=75). There were no statistically significant differences in age between groups by Kruskal-Wallis with Dunn’s test for multiple comparisons. There were also no differences in distributions of male/female sex between groups by Fisher’s exact test. There were significant differences in *APOE*4+ allele status between groups (Table I).

Indicators of inflammation from peripheral blood and CSF were assessed in a subset of participants with available measures. Participants with AD had significantly higher white blood cell counts compared to MCI (p=0.0239) and significantly lower basophil percentages compared to CN (p=0.0309). Participants with MCI had significantly lower erythrocyte sedimentation rates compared to CN (p=0.0188). Participants did not vary by percentages of neutrophils, lymphocytes, monocytes, levels of C-reactive protein, or blood brain barrier integrity (ratio of CSF albumin/serum albumin).

A subgroup of the cohort with CSF measurements of Aβ42/40, p-Tau 181, and total Tau were characterized into ATN subgroups (CN n=27, MCI n=27, AD n=65). Significant differences were observed between all three disease groups’ levels of Aβ42/40 and levels of p-Tau. Additionally, both CN and MCI groups had significantly lower levels of total Tau compared to the AD group (Table I). In order to better understand the differences in groups defined primarily by signs and symptoms (disease status) compared to CSF AD-related biomarkers (ATN status), we also compared groups according to their ATN status. Characteristics of the study populations were compared between groups defined by the ATN Continuum, AD portion: A-T-N- (n=31), A+T-N- (n=17), A+T+N- (n=37), and A+T+N+ (n=39). People in other ATN groups (n=5), such as A-T-N+ were not included in the analysis. There were n=71 participants who could not be defined by ATN group because of the lack of CSF material available. There were no statistically significant differences in age (by Mann-Whitney tests) or gender (by Fisher’s exact test) between groups (Supplementary Table I). Upon stratification by ATN category, increased neutrophil counts in A+T-N- compared to A-T-N-, and higher WBC counts in A+T-N- compared to both A-T-N- and A+T+N- was observed (Supplementary Table I).

### Levels of sTREM2 and multiple other inflammatory factors tend to be lower in AD

We measured levels of plasma sTREM2 and 38 cytokines and compared disease and ATN groups. Levels of plasma sTREM2 were significantly lower in the AD group compared to the MCI group (p=0.0421, Figure 1A). We found that levels of 6 plasma cytokines had significant differences between disease groups. Levels of fractalkine, IL-5, and EGF were also higher in MCI compared to AD (fractalkine p=0.0029; IL-5 p=0.0134; EGF p=0.0434). We observed higher levels of IL-1RA in CN compared to MCI (p=0.0359), and higher levels of Flt-3L in CN compared to AD (p=0.0137). Finally, levels of IL-17A were higher in MCI compared to both CN (p=0.0164) and AD (p=0.0436). Tests were performed using Kruskal-Wallis adjusted for multiple comparisons after the removal of outliers (Figure 1B-D). Supplementary Table II indicates numbers of outliers removed per group per analyte.

**Figure 1:**
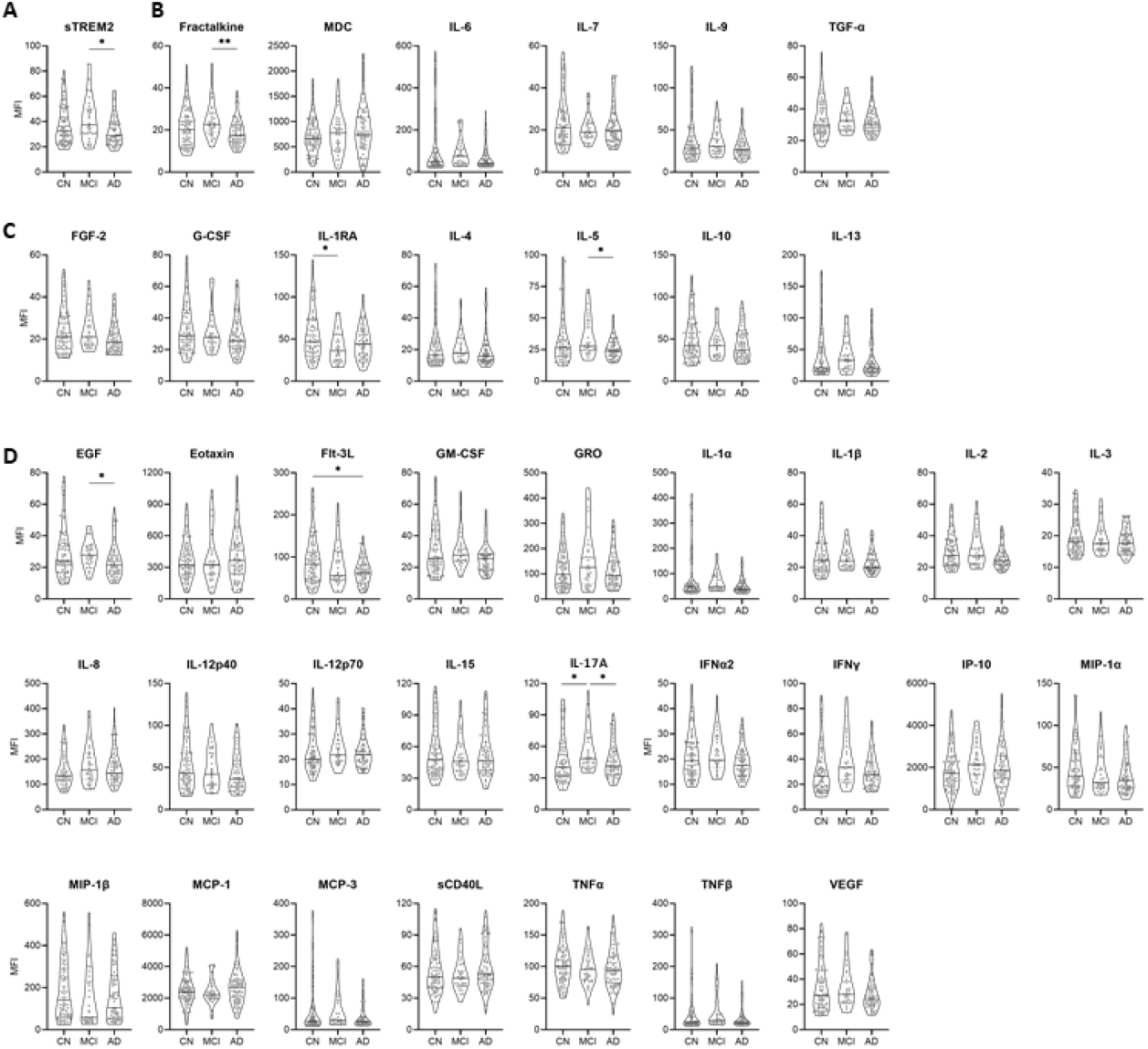
Levels of sTREM2 and multiple other inflammatory factors tend to be lower in AD. sTREM2 was significantly lower in AD, compared to MCI. (A) The immunoregulatoy/pleiotropic factor, fractalkine, was significantly lower in AD, compared to MCI. (B) The anti-inflammatory factor IL-1RA was significantly lower in MCI, compared to controls. IL-5 was significantly lower in AD, compared to MCI. (C) The pro-inflammatory factor EGF was significantly lower in AD, compared to MCI. Flt-3L was significantly lower in AD, compared to controls. IL-17A was significantly higher in MCI compared to both controls and AD. (D) Levels were compared between disease groups by Kruskal-Wallis tests with Dunn’s test for multiple comparisons after removal of outliers. Supplementary Table II shows numbers of outliers removed per disease group for each cytokine using the ROUT method. *p<0.05; **p<0.01; ***p<0.001; ****p<0.0001

### Levels of immunoregulatory/pleiotropic factors tend to be higher in A+T-N- while anti-inflammatory and pro-inflammatory factors tend to be higher in A+T+N-

We similarly compared levels of sTREM2 and cytokines between the ATN group sub-cohort. In order to focus on designations related to AD progression (8), we analyzed only participants defined as A-T-N-, A+T-N-, A+T+N-, or A+T+N+ (124 participants out of 200 in the total cohort). Groups were compared along the ATN continuum by ManN- Whitney tests after outlier removal (A-T-N- vs A+T-N-, A+T-N- vs A+T+N-, and A+T+N- vs A+T+N+). Levels of sTREM2 did not differ between ATN groups (Figure 2A). Levels of immunoregulatory/pleiotropic cytokines fractalkine and IL-7 were higher in the A+T-N- group compared to the A-T-N- and A+T+N- groups (fractalkine p=0.0196 and 0.0017, IL-7 p=0.0156 and 0.0031, respectively) (Figure 2B). The A+T+N+ group had significantly lower levels of anti-inflammatory cytokines FGF-2 and IL-5 compared to the A+T+N- group (FGF-2 p=0.0237, IL-5 p=0.0395) (Figure 2C). Similarly, levels of pro-inflammatory cytokines GRO, IL-1β, IL-2, IL-3, IL-8, IL-15, IFNγ, MCP-3, and sCD40L were lower in A+T+N+ compared to A+T+N-, in addition to higher levels of MCP-3 in A+T-N- compared to A-T-N- (Figure 2D). These analyses were completed after outlier removal; outliers removed per group per analyte are shown in Supplementary Table III.

**Figure 2:**
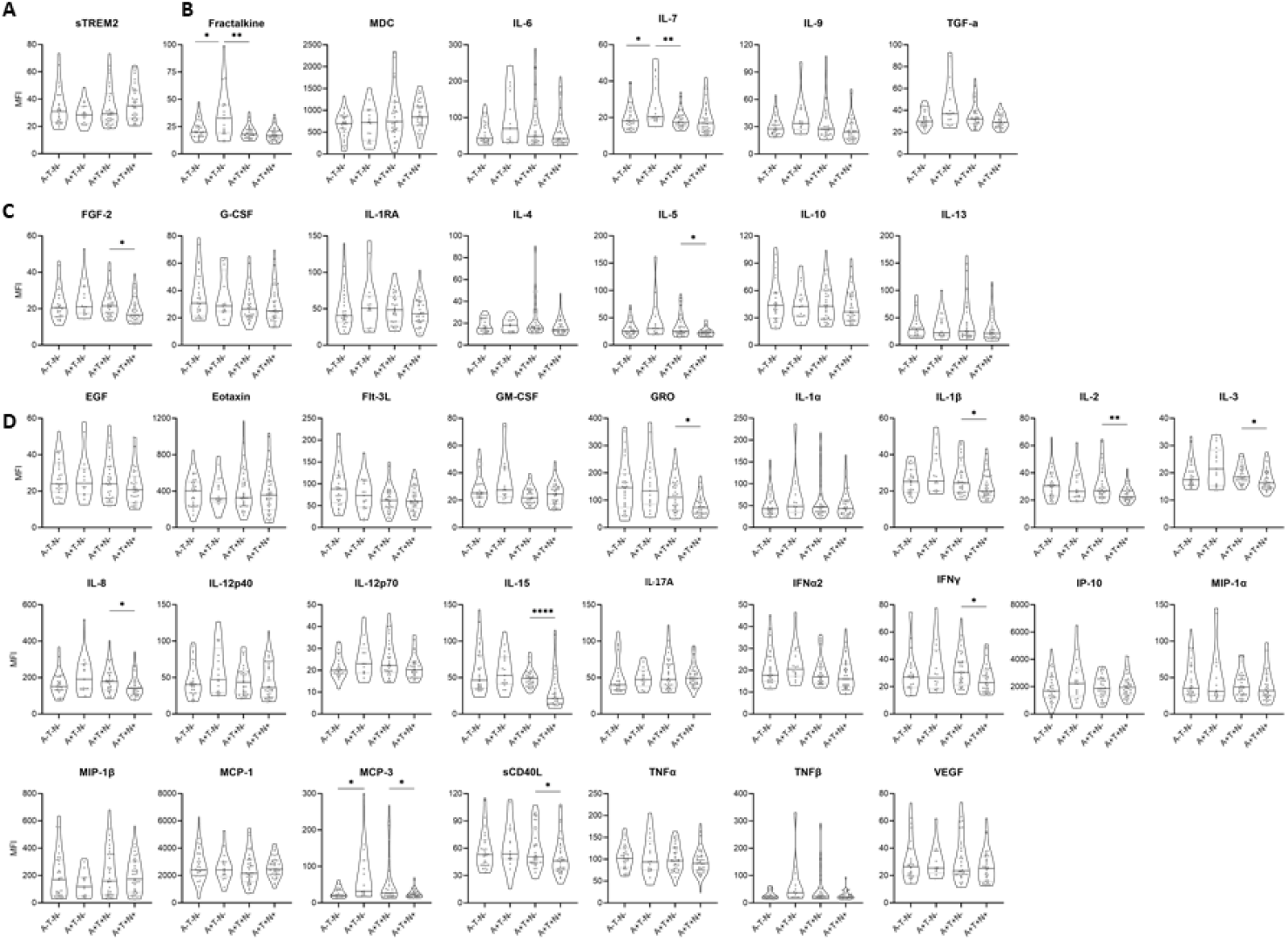
Levels of immunoregulatory/pleiotropic factors tend to be higher in A+T-N- while anti-inflammatory and pro-inflammatory factors tend to be higher in A+T+N-. Levels of sTREM2 were not different between groups. (A) The immunoregulatory/pleiotropic factor, fractalkine, was significantly higher in the A+T-N-, compared to A-T-N- and A+T+N. IL-7 was significantly higher in the A+T-N-group, compared to A-T-N- and A+T+N-. (B) The anti-inflammatory factors, FGF-2 and IL-5 were significantly higher in A+T+N-, compared to A+T+N+. (C) The pro-inflammatory factors GRO, IL-1β, IL-2, IL-3, IL-8, IL-15, IFNγ, MCP-3 and sCD40L are lower in A+T+N+, compared to A+T+N-. (D) Levels were compared by Mann-Whitney tests after removal of outliers. Groups were compared as follows: A-T-N- vs A+T-N-; A+T-N- vs A+T+N-, and A+T+N- vs A+T+N+. Supplementary Table III shows numbers of outliers removed per disease group for each cytokine using the ROUT method. *p<0.05; **p<0.01; ***p<0.001; ****p<0.0001

### MCI and AD showed significantly diminished correlations of sTREM2 and inflammatory factors

We performed Spearman correlations on all 39 markers (sTREM2 and 38 cytokines) compared to all other markers within each group to better understand the relationships in the broader immune response (Figure 3. Hierarchical clustering was performed to group sTREM2 and cytokines with similar patterns, in order to better understand if sTREM2 was grouping with anti-inflammatory, pro-inflammatory, or pleiotropic/immunoregulatory cytokines.

**Figure 3:**
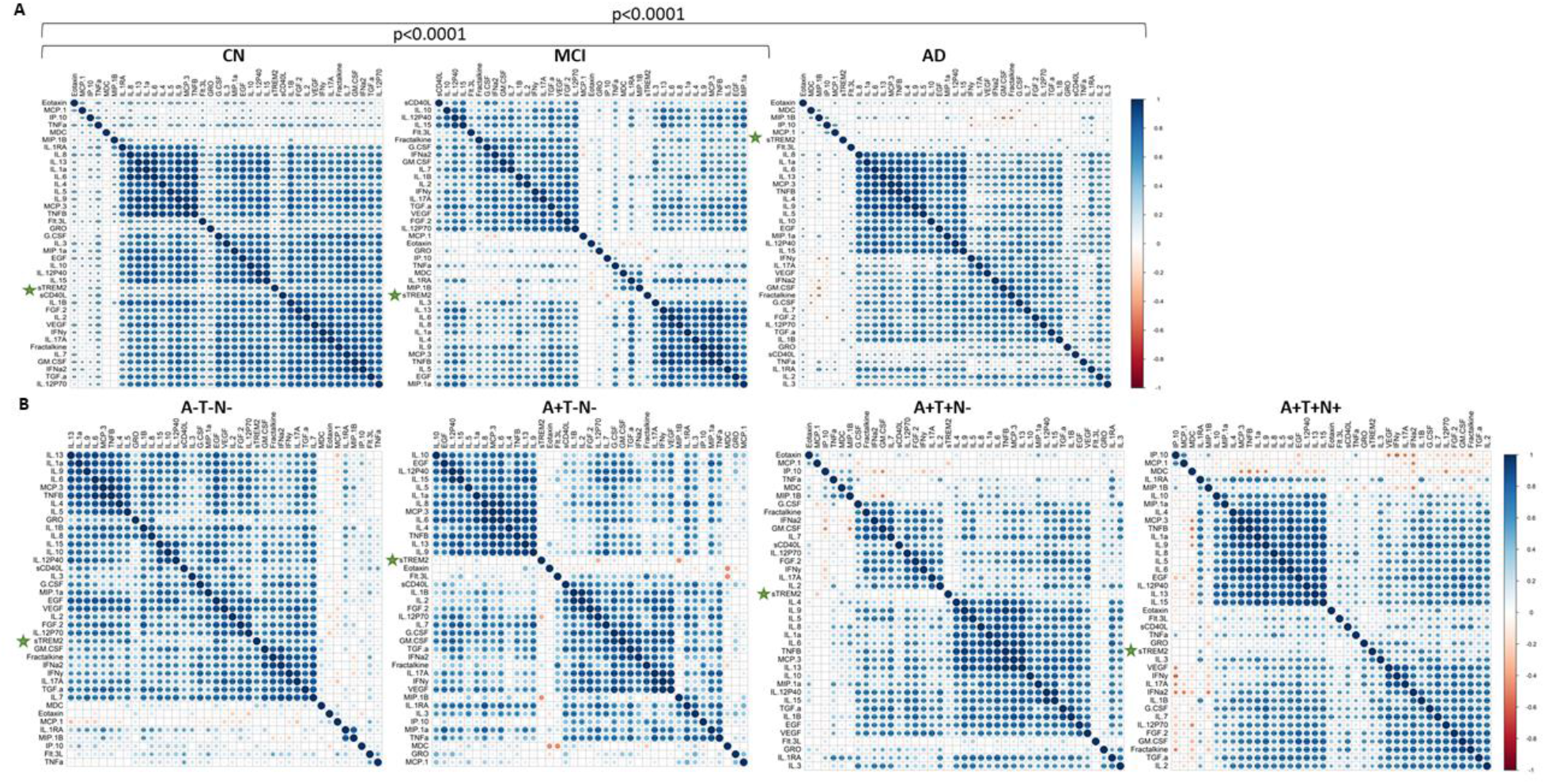
MCI and AD showed significantly diminished correlations of sTREM2 and inflammatory factors. sTREM2 and each cytokine were correlated to each other in (A) disease groups and (B) ATN groups. Clustering was performed to show similar patterns of correlations within groups. Steiger tests to compare matrices were run and significant differences are noted. Correlation directions are shown by a color gradient; positive correlations are shown in blue and negative correlations are red. Significant correlations are noted by displaying asterisks for p values. *p<0.05; **p<0.01; ***p<0.001; ****p<0.0001

Outlier removal was not possible in this comparison due to limitations in the software package; introducing missing values resulted in the elimination of participants, so all data points were included. In the correlation plots of disease groups, CN showed significant positive correlations of sTREM2 with 34 of the 38 inflammatory factors (the exceptions were eotaxin, MDC, MIP-1β, and MCP-1) (Figure 3A). This was in contrast to the MCI group, in which sTREM2 significantly correlated with only IL-1α, IL-1β, IL-10, and EGF (Figure 3A). In the AD group, sTREM2 significantly correlated with 12 of the 38 inflammatory factors (Figure 3A), three of which were in common with those seen in MCI. Additionally, we used the Steiger test to compare correlation matrices and found that the CN group was significantly different than both the MCI and AD groups (p<0.0001). In a separate analysis, sTREM2 was correlated to each cytokine after outlier removal. We found that 17 cytokines were significantly positively correlated with sTREM2 levels in the CN group (EGF, FGF-2, TGFα, GM-CSF, fractalkine, IFNα2, IFNγ, IL-10, sCD40L, IL-17A, IL-2, IL-7, IP-10, MIP-1α, TNFα, and VEGF), while zero were significantly correlated in the MCI group and five were significantly positively correlated in the AD group (EGF, IL-10, IL-1β, IL-3, and VEGF) (Supplementary Table IV).

The ATN groups showed similar patterns of correlation matrices as the disease groups. Similar to the CN group, A-T-N- had significant positive correlations of sTREM2 with of the cytokines, including IL-13, IL-1α, IL-9, IL-6, MCP-3, TNFβ, IL-4, IL-5, GRO, IL-1β, IL-8, IL-10, IL-12p40, EGF, VEGF, IL-2, FGF-2, IL-12p70, GM-CSF, fractalkine, IFNα2, IFNγ, IL-17A, TGFα, and IL-7. A+T-N- showed no significant correlations of sTREM2 with cytokines. A+T+N- showed significant correlation of sTREM2 with IL-4, IL-9, IL-1α, IL-6, TNFβ, MCP-3, IL-13, IL-10, IL-1β, and EGF, and A+T+N+ showed positive correlations of sTREM2 with IL-1RA, TNFβ, IL-6, EGF, GRO, IL-3, VEGF, IFNγ, IL-17A, IFNα2, IL-1β, G-CSF, IL-7, IL-12p70, FGF-2, and GM-CSF (Figure 3B). The groups had had different patterns of hierarchical clustering, but the correlation matrices were not significantly different by the Steiger test (Figure 3B). In a separate analysis, Spearman correlations were performed after the removal of outliers. There were fewer significant correlations after outlier removal. In the A-T-N- group, sTREM2 correlated only with IL-5, MDC, EGF, and IP-10; in A+T-N- sTREM2 correlated only with IL-4 and IL-12p70; in A+T+N-, sTREM2 correlated only with IL-10, TGFα, EGF, and IL-1β; in A+T+N+, sTREM2 correlated with FGF-2, IL-7, GM-CSF, GRO, IFNα2, IL-1β, and IL-3 (Supplementary Table IV).

### sTREM2 showed more significant relationships with inflammatory factors in CN than MCI and AD

Single linear regressions were run to measure the effect of changes in sTREM2 (the independent variable) on each of the 38 cytokines (as the dependent variables). The CN group showed more significant relationships than the MCI and AD groups; TGFα, FGF-2, G-CSF, GM-CSF, IFNα2, IFNγ, IL-2, and MIP-1α showed significant Beta coefficients only in the CN group, while fractalkine was significant in both the CN and MCI groups, IL-7, and EGF were significant in the CN and AD groups, IL-1β and IL-3 were significant in only AD, and IL-10 was significant in all three disease groups (Supplementary Table V).

### sTREM2 networks are altered at the MCI or A+T-N- stages

Network analysis of the correlation between sTREM2 and inflammatory factors revealed substantial differences in the sTREM2 modules between groups. The CN group showed a dense module with sTREM2 at the center of the hub, connected to many cytokines, while the MCI group showed a very sparse network where only three cytokines connected to sTREM2, while the AD group showed a small network connected to sTREM2, with a higher degree of connectivity between other inflammatory factors (Figure 4A). The MCI and AD groups also showed networks of inflammatory factors that were not connected to sTREM2, while the CN group did not. Similarly, the A-T-N- network analysis showed a strong module around sTREM2, which was not seen in A+T-N- (no connections with sTREM2), A+T+N-, or A+T+N+, which both showed sparser networks with hubs outside of sTREM2 (Figure 4B).

**Figure 4:**
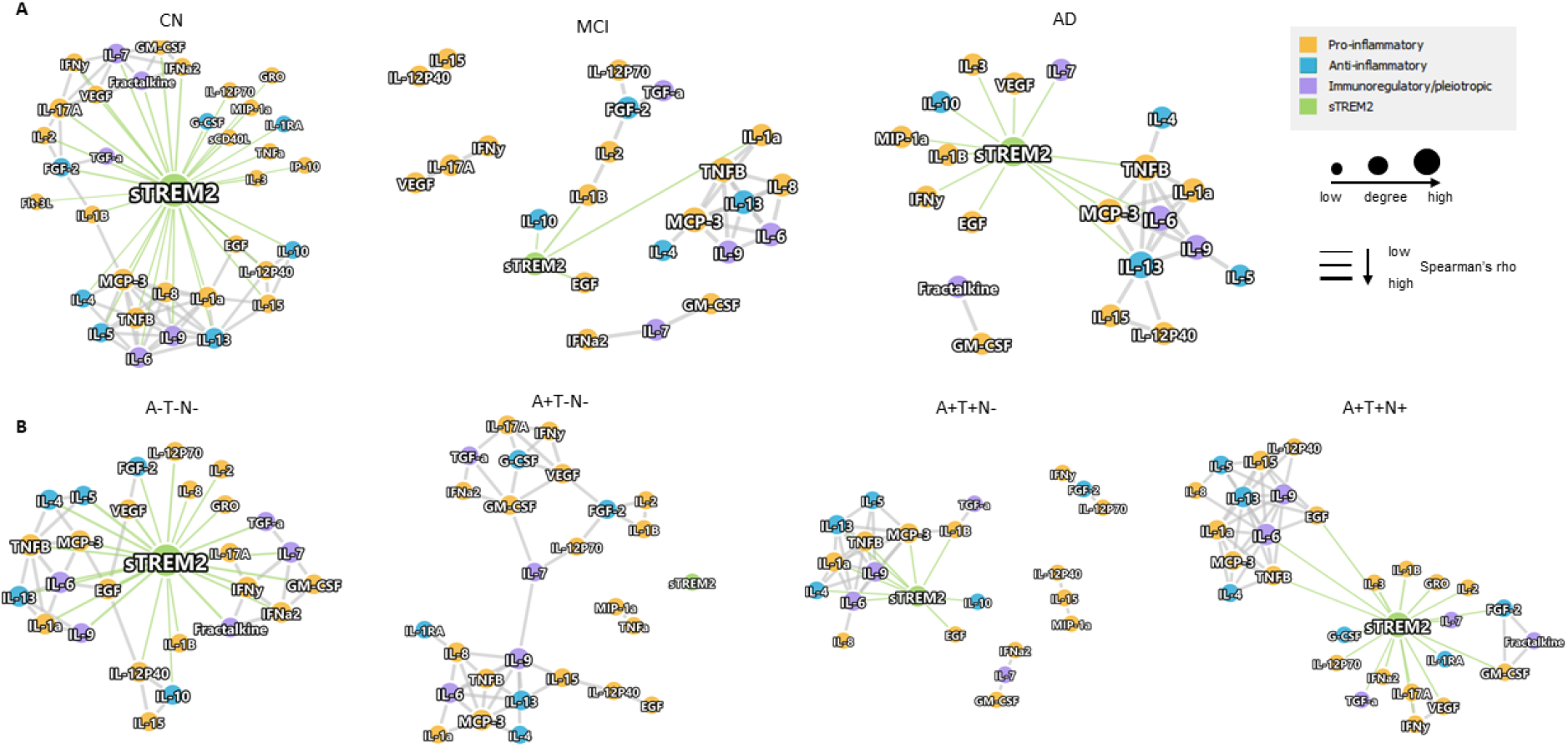
Plasma sTREM2 networks are altered at the MCI and A+T-N- stages. Network analyses were performed to visualize Spearman rho correlations between sTREM2 and inflammatory factors. Within disease groups, CN showed a strong sTREM2 hub with connections to many inflammatory factors, while there were few connections in MCI and AD (A). Similarly, A-T-N- showed a large sTREM2 module, which was absent in A+T-N- and diminished in A+T+N- and A+T+N+ (B). Inflammatory factor nodes are colored according to their broad functional categories: pro-inflammatory (orange, anti-inflammatory (blue), immunoregulatory/ pleiotropic (purple), and sTREM2 is green. Node size and thickness of lines are related to levels of Spearman rho. Inclusion of inflammatory factors was based on Spearman r>0.8 and p<0.05, and sTREM2 correlations were shown with r>0 and p<0.05.

### Regardless of disease status inflammatory factors tend to be higher in the high sTREM2 tertile

Participants were separated by sTREM2 tertile by ordering sTREM2 MFI levels smallest to largest and evenly dividing them into three groups, regardless of disease status. Tertile 1 refers to the lowest sTREM2 levels and Tertile 3 refers to the highest sTREM2 levels. Bar graphs show numbers of each disease group per tertile (Figure 5). There were significantly higher proportions of CN in Tertile 3 vs MCI and MCI in Tertile 3 vs AD, but not CN vs AD by Fisher’s exact test, indicating CN and MCI represented a majority of the high sTREM2 values. When we compared distributions of disease groups within sTREM2 tertiles, half of the CN group was in tertile 3 and the other half in tertile 1. In contrast, a greater proportion of MCI were in tertile 3 compared to tertile 1 and fewer AD were in Tertile 3 compared to tertile 1. Similarly, when all three disease groups were compared there was a significant difference by Chi square test (Figure 5). Tertile 1 showed a range in sTREM2 MFI of 16.75-28.25, and Tertile 3 had a range of 43.0-108.0 MFI (Figure 6). We compared levels of cytokines between Tertiles 1 and 3 and found significant differences in 31 of the 39 inflammatory factors measured. In each significant difference, the group with high sTREM2 (Tertile 3) also had higher levels of cytokines. The cytokines that were significantly higher in Tertile 3 were immuneregulatory/ pleiotropic cytokines fractalkine, IL-6, IL-7,IL-9, and TGFα (Figure 6), anti-inflammatory cytokines FGF-2, G-CSF, IL-4, IL-5, IL-10, and IL-13 (Figure 5C) and pro-inflammatory cytokines EGF, GM-CSF, GRO, IL-1α, IL-1β, IL-2, IL-3, IL-8, IL-12p40, IL-12p70, IL-15, IL-17A, IFNα2, IFNγ, IP-10, MIP-1α, MCP-3, sCD40L, TNFα, TNFβ, and VEGF (Figure 6D). Interestingly, there were no cytokines measured with higher levels in Tertile 1 compared to Tertile 3. Outliers were removed prior to this analysis; numbers of outliers-removed per analyte per group are shown in Supplementary Table VI.

**Figure 5:**
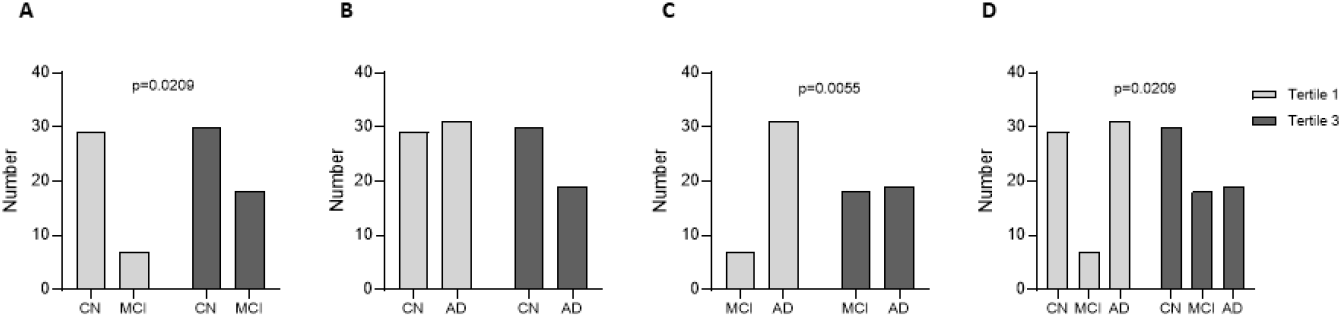
AD participants were disproportionally in the lowest sTREM2 tertile, while MCI disproportionally was in the highest sTREM2 tertile. CN, MCI, and AD participants were separated into tertiles based on low (Tertile 1) and high (Tertile 3) levels of sTREM2. There were significantly different proportions of CN vs MCI (A), and MCI vs AD (C), but not CN vs AD (B) by Fisher’s exact test. Similarly, when all three disease groups were compared there was a significant difference by Chi square test (D).

**Figure 6:**
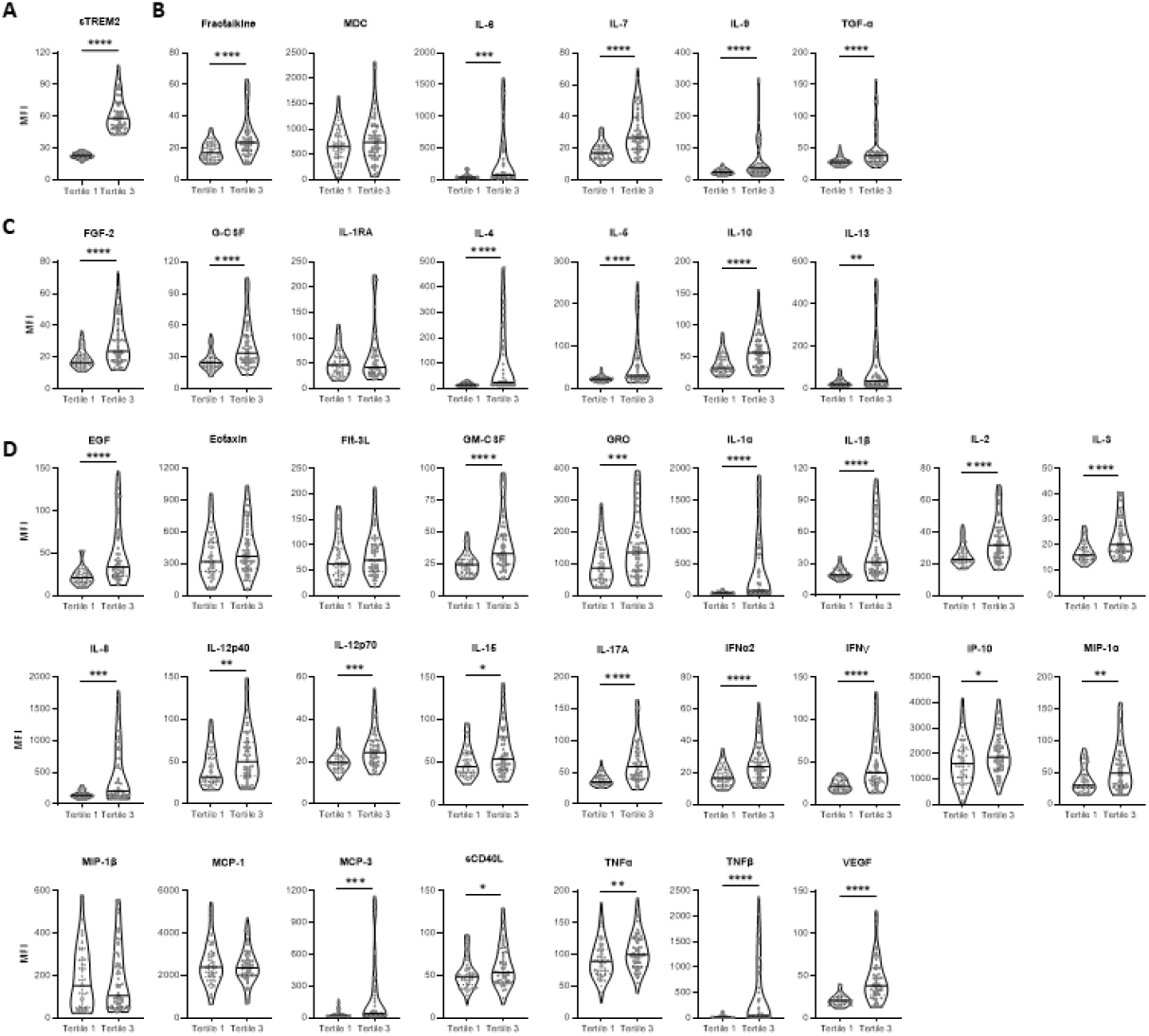
Inflammatory factors tend to be higher in the high sTREM2 tertile. sTREM2 Tertile 1 showed a range in sTREM2 MFI of 16.75-28.25, and Tertile 3 had a range of 43.0-108.0 MFI. (A) The immunoregulatory/ pleiotropic factors that were significantly higher in the high sTREM2 tertile included: fractalkine, IL-6, IL-7, IL-9, ad TGF-α. (B) The anti-inflammatory factors that were significantly higher in the high sTREM2 tertile included: FGF-2, G-CSF, IL-4, IL-5, IL-10, and IL-13. (C) The pro-inflammatory factors that were significantly higher in the high sTREM2 tertile included: EGF, GM-CSF, GRO, IL-1α, IL-1β, IL-2, IL-3, IL-8, IL-12p40, IL-12p70, IL-15, IL-17A, IFNα2, IFNγ, IP-10, MIP-1α, MCP-3, sCD40L, TNFα, TNFβ, and VEGF (D). Levels were compared between sTREM2 Tertile 1 (low) and sTREM2 Tertile 1 (high) by Mann-Whitney tests after removal of outliers. Supplementary Table V shows numbers of outliers removed per disease group for each cytokine using the ROUT method. *p<0.05; **p<0.01; ***p<0.001; ****p<0.0001

### Inflammatory factor correlations were significantly altered with low sTREM2

Correlation matrices plots were generated in order to show the broader inflammatory activity differences between sTREM2 Tertile 1 (low) (Figure 7A) and Tertile 3 (high) (Figure 7B). The plots shown are without outlier removal. The patterns of correlation were very different between Tertile 1 and Tertile 3, shown visually by a color gradient of positive (blue) to negative (red) correlations, with white showing no correlation (Spearman r=0). Overall, Tertile 1 had fewer significant correlations than Tertile 3 for both sTREM2 with the cytokines and between the cytokines. Correlation matrices were compared by the Steiger test and found to be significantly different (p<0.0001). In a separate analysis that allowed for the removal of outliers, sTREM2 was correlated with each cytokine within tertiles. Tertile 1 (low sTREM2) did not significantly correlate with any of the 38 cytokines, while Tertile 3 (high sTREM2) had significantly positive correlations with EGF, FGF-2, G-CSF, GM-CSF, fractalkine, IFNα2, IFNγ, IL-12p40, IL-15, sCD40L, IL-17A, IL-1α, IL-9, IL-1β, IL-2, IL-3, IL-4, IL-5, IL-6, IL-7, IL-8, MIP-1α, TNFβ, and VEGF (Spearman correlation after outlier removal shown in Supplementary Table VII).

**Figure 7:**
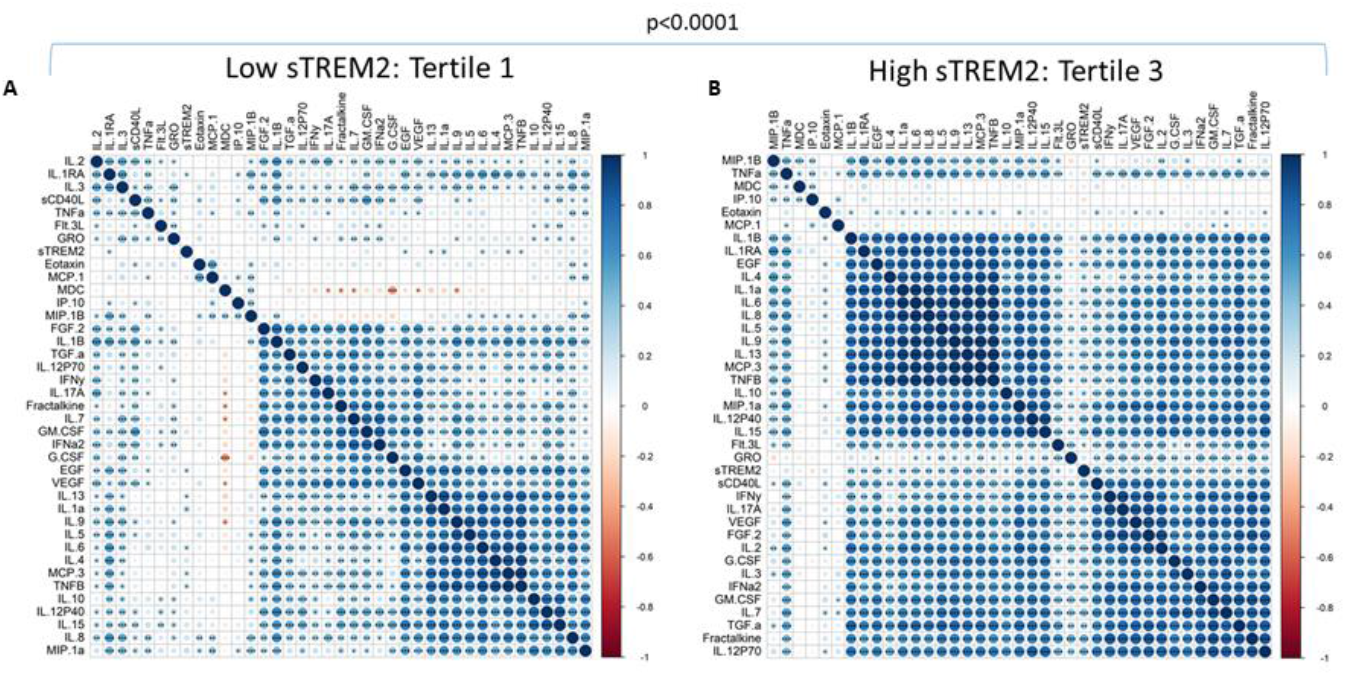
Inflammatory factor correlations were significantly altered with low sTREM2. sTREM2 and each cytokine were correlated to each other in (A) sTREM2 Tertile 1 and (B) sTREM2 Tertile 3. Clustering was performed to show similar patterns of correlations within groups. Steiger tests to compare matrices were run and significant differences are noted. Correlation directions are shown by a color gradient; positive correlations are shown in blue and negative correlations are red. Significant correlations are noted by displaying asterisks for p values. *p<0.05; **p<0.01; ***p<0.001; ****p<0.0001

## Discussion

The purpose of this study was to explore the relationship between plasma sTREM2 and inflammatory activity in AD. We sought to accomplish this by comparing plasma sTREM2 and plasma inflammatory factors in clinical or ATN categories.

An association between sTREM2 and inflammatory factors was observed that was different in the CN and A-T-N- groups compared to the other groups defined by AD symptoms (MCI and AD) or AD-related pathology (A+T-N-, A+T+N-, and A+T+N+). Overall, patterns of inflammatory activity and the relationship with sTREM2 was drastically altered in early stage AD, and remains altered in the later-stages of AD, although less so, suggesting that TREM2 related inflammatory activity plays a critical role early in disease progression especially at the MCI and A+T-N- stage. Together, these results are supported by a previous report that describes higher CSF sTREM2 as attenuating *APOE*4-related risk for cognitive decline and neurodegeneration suggesting that indeed early elevation of sTREM2 plays a critical role in AD progression (36). In addition, studies in mouse models suggest that TREM2 function impedes tau seeding in neuritic plaques (37) and potentially restrains the enhancement of tau accumulation and neurodegeneration initiated by β-amyloid pathology (38). Together, this suggests that in the periphery there are initial early stage alterations in TREM2 related inflammatory activity that corresponds to a broad alteration in the inflammatory landscape and is followed by less diminished alterations in sTREM2 related inflammatory activity in the later stages of the disease.

Interestingly, there were instances in which none of the disease groups showed sTREM2 correlations with inflammatory factors. The four inflammatory factors that did not correlate significantly with sTREM2 in the CN group (eotaxin, MDC, MIP-1β, and MCP-1), did not significantly correlate in the MCI and AD group either, and were not different between sTREM2 tertiles suggesting that those inflammatory factors are not associated with sTREM2 in this cohort even though previous evidence suggests they are altered in AD (39, 40).

Several cytokines stood out as having the most significant findings. Between groups, fractalkine was significantly higher in MCI than AD, and significantly higher in A+T-N- compared to both A-T-N- and A+T+N-. Fractalkine was significantly correlated with sTREM2 only in the CN and sTREM2 Tertile 3 groups (after outlier removal), and was significantly elevated in sTREM2 Tertile 3. Fractalkine is an immunoregulatory cytokine which has been shown to reduce inflammatory signaling in activated microglia and reduce tau pathology when over-expressed (41, 42) and has been implicated in other studies (43–46). Additionally, both fractalkine and TREM2 are cleaved by the same enzyme, a disintegrin and metalloproteinase domain-containing protein 10 (ADAM-10), which may link the downstream function of the two proteins (18, 19, 47). IL-5 was also higher in MCI than AD, higher in A+T+N- compared to A+T+N+, and higher in sTREM2 Tertile 3 compared to Tertile 1. IL-5 has been shown to be neurotrophic (48) and, in cell culture, increase cell survival and inhibit phosphorylation of tau (49) and has been previously shown to be diminished in AD (50). IL-1α and IL-1β were both higher in A+T-N- than A-T-N- and higher in sTREM2 Tertile 3 than Tertile 1. IL-1α was also higher in A+T-N- than A+T+N-. Although not significantly different between disease groups, IL-1β correlated with sTREM2 only in the AD, A+T+N-, and A+T+N+ groups, after outlier removal. IL-1α and IL-1β are known pro-inflammatory cytokines and previous reports have shown increased levels of both cytokines in serum from patients with AD (51); however, IL-1β overexpression was found to be associated with reduced amyloid plaques and levels of Aβ42 and Aβ40 in a mouse model of AD (52) and IL-1β was shown to down-regulate expression of TREM2 mRNA in cultures of human peripheral blood monocytes and synovial fluid macrophages from patients with rheumatoid arthritis (53). Finally, IL-17A was significantly higher in MCI compared to both CN and AD, and significantly higher in sTREM2 Tertile 3 compared to Tertile 1. IL-17A was significantly correlated with sTREM2 in the CN and A-T-N- groups, and showed a significant relationship with sTREM2 by linear regression only in the CN group (after outlier removal). IL-17A is a strong pro-inflammatory cytokine that signals to microglia, has been linked to systemic inflammatory diseases including autoimmune and metabolic diseases and AD-related inflammation in the brain (54–56).

Comparisons of sTREM2 tertiles, which were done regardless of disease or ATN status, give evidence of sTREM2-related inflammatory activity. Tertile 3 (high sTREM2) had significantly higher levels of 31 cytokines while tertile 1 did not have higher levels of any cytokine. Given that inflammatory activity matrices were significantly different between tertiles and MCI is disproportionally in tertile 3 while AD is in tertile 1, these results support the idea that TREM2 related inflammatory activity plays a key role in early stage AD while this role changes later in the disease.

A limitation of this exploratory study is small sample size, especially for the ATN groups. Although we did gather evidence for fractalkine, IL-1α IL-1β, IL-5, and IL-17A, our findings will need to be tested in a larger cohort to validate the correlations observed. Even though, a preliminary power analysis suggested a sample size of 25 per group would achieve the desired power, the data should be approached with caution since there is a potential for false negative results for some of the analytes with detectable levels in the very low range. Another limitation is that some important inflammatory factors may have been missed since the thirty-eight inflammatory factors available on the inflammatory panel utilized is not comprehensive of the multiple immune factors present in the circulation. Additionally, another limitation is that we utilized Aβ42/Aβ40 for “A”, p-Tau for “T” and t-Tau for “N” as in other reports (57). Others have argued that only “A” and “T” groups should be used or that imaging or neurofilament light should be used for “N” (58, 59). As with any test that uses a cutoff value for positivity, there is likely a grey area of overlap between the groups that could not be completely separated. Furthermore, given the identified alterations in basophil levels and erythrocyte sedimentation rate in AD the results should be approached with caution since, immune factor levels could have been altered due to factors such as undiagnosed inflammatory illnesses or impending infections. Future studies will benefit from increased sample size, more diverse inflammatory panels and single cell-based analyses to further understand the underlying source of these results. Importantly, longitudinal studies will help tease apart these inflammatory effects on disease progression.

In this exploratory study, we identified patterns of inflammatory factor relationships with plasma sTREM2 that differed by disease group, ATN group, and sTREM2 tertile. The cytokines that were found to have the strongest relationship with sTREM2 across comparison groups were fractalkine, IL-5, IL-1α, IL-1β, and IL-17A. Plasma sTREM2 was linked to inflammatory activity in the peripheral circulation, in that strong connections and patterns observed in the groups without AD symptoms or CSF biomarkers, CN and A-T-N-, were profoundly altered in the A+T-N- and MCI stages. Together, this suggests that the pathological amyloid stage, prior to the pathological tau stage, may be a critical stage for peripheral immune system intervention in AD pathogenesis. These findings lay the groundwork for research and therapeutic strategies that seek to understand or target the immune system in AD.

## List of abbreviations

AD: Alzheimer’s disease
TREM2: triggering receptor expressed on myeloid cells 2
sTREM2: soluble triggering receptor expressed on myeloid cells 2
ATN: <u>a</u>myloid beta, phosphorylated <u>t</u>au, and <u>n</u>eurodegeneration
Aβ: amyloid beta
CSF: cerebrospinal fluid
CN: cognitively normal
MCI: mild cognitive impairment
p-Tau: phosphorylated tau
t-Tau: total tau
LRCBH-Biobank: Cleveland Clinic Lou Ruvo Center for Brain Health Aging and Neurodegenerative Disease Biobank
CADRC: Cleveland Alzheimer’s Disease Research Center
CBC: Complete blood counts
APOE: apolipoprotein E
EGF: Epidermal Growth Factor
FGF-2: Fibroblast Growth Factor 2
TGF-α: Transforming Growth Factor alpha
G-CSF: Granulocyte-colony stimulating factor
Flt-3L: FMS-like tyrosine kinase 3 ligand
GM-CSF: Granulocyte-Macrophage Colony Stimulating Factor
IFN: interferon
GRO: growth-regulated oncogene
IL: interleukin
MCP: Monocyte chemotactic protein
MDC: Macrophage-derived chemokine
sCD40L: soluble CD40-ligand
IL-1RA: interleukin 1 receptor agonist
IP-10: interferon-gamma inducible protein
MIP: macrophage inflammatory protein
TNF: tumor necrosis factor
VEGF: Vascular Endothelial Growth Factor
MFI: Mean fluorescence intensity

## Declarations

Ethics approval and consent to participate: Participants consented under the Lou Ruvo Center for Brain Health Aging and Neurodegenerative Disease Biobank (LRCBH-Biobank) and the Cleveland Alzheimer’s Disease Research Center (CADRC) protocols approved by the Cleveland Clinic Institutional Review Board.

## Authors’ contributions

GEW analyzed and interpreted data, prepared figures, and wrote the manuscript text. MK performed assays and prepared data. EDT analyzed data and prepared figures. YS developed assays. JP and SR characterized participants’ clinical status and performed neurological testing. HF, YZ, and FC contributed analyses, figure development, and manuscript editing. TMD and SS contributed result interpretation and manuscript development. JBL performed neurological testing, participant consensus, data interpretation and manuscript development. LMB oversaw the study design, assays completion, analysis, figure development, and manuscript preparation. All authors read and approved the final manuscript.

---

**Supplementary Table I:** Cohort demographics within ATN groups. A subset of participants were classified into ATN groups. Characteristics of the study populations were compared between groups defined by the ATN Continuum, AD portion: A-T-N- (n=31), A+T-N- (n=17), A+T+N- (n=37), and A+T+N+ (n=39). People in other ATN groups (n=5), such as A-T-N+ were not included in the analysis. There were n=71 participants who could not be defined by ATN group because of the lack of CSF material available. There were no statistically significant differences in age (by Mann-Whitney tests) or gender (by Fisher’s exact test) between groups. Significant differences are noted in bold.

| Group | A-T-N- | A+T-N- | A+T+N- | A+T+N+ | A-T-N-<br>vs<br>A+T-N- | A+T-N-<br>vs<br>A+T+N- | A+T+N-<br>vs<br>A+T+N+ |
| --- | --- | --- | --- | --- | --- | --- | --- |
| **n** | 31 | 17 | 37 | 39 |  |  |  |
| **Age range (years)** | 55-79 | 56-76 | 55-84 | 56-86 |  |  |  |
| **Average age (years)** | 64.4 | 66.8 | 66.7 | 67.1 | 0.1167 | 0.7924 | 0.9320 |
| **% Male** | 64.5 | 70.6 | 54.1 | 38.5 | 0.7566 | 0.3723 | 0.3518 |
| **% APOE4+** | 45.2 | 76.5 | 75.7 | 61.5 | 0.0666 | >0.9999 | 0.2224 |
| **Clinical data sub-cohort** |  |  |  |  |  |  |  |
| **CBC** | n=24 | n=15 | n=34 | n=32 |  |  |  |
| WBC mean (SD) | 5.20 (1.23) | 6.35 (1.28) | 5.59 (1.68) | 5.33 (1.24) | **0.0064** | **0.0358** | 0.8783 |
| Absolute Neutrophil Count mean (SD) | 3.22 (0.99) | 4.82 (2.24) | 3.37 (0.88) | 3.52 (1.20) | **0.0140** | 0.0660 | 0.7996 |
| Neutrophil % mean (SD) | 59.25 (10.09) | 64.35 (8.354) | 63.58 (7.98) | 60.91 (10.43) | 0.1018 | 0.5730 | 0.5257 |
| Absolute Lymphocyte Count mean (SD) | 1.44 (0.40) | 1.55 (0.42) | 1.32 (0.46) | 1.37 (0.46) | 0.5204 | 0.2002 | 0.8953 |
| Lymphocyte % mean (SD) | 28.2 (7.61) | 24.49 (8.36) | 25.39 (7.29) | 27.59 (9.55) | 0.1366 | 0.3577 | 0.5857 |
| Absolute Monocyte Count mean (SD) | 0.46 (0.11) | 0.50 (0.22) | 0.47 (0.12) | 0.46 (0.10) | 0.9792 | 0.8518 | >0.9999 |
| Monocyte % mean (SD) | 8.56 (1.17) | 7.94 (1.18) | 7.97 (1.95) | 8.92 (1.84) | 0.1120 | 0.8342 | 0.0737 |
| Absolute Basophil Count mean (SD) | 0.022 (0.014) | 0.028 (0.012) | 0.023 (0.014) | 0.022 (0.012) | 0.1378 | 0.2307 | 0.8100 |
| Basophil % mean (SD) | 0.50 (0.32) | 0.38 (0.23) | 0.42 (0.23) | 0.43 (0.21) | 0.2565 | 0.5845 | 0.8543 |
| **Sed rate** | n=21 | n=15 | n=34 | n=32 |  |  |  |
| Mean (SD) | 7.7 (4.68) | 5.8 (2.18) | 8.24 (4.89) | 7.3 (3.75) | 0.1842 | 0.0986 | 0.5796 |
| **CRP** | n=23 | n=13 | n=31 | n=32 |  |  |  |
| Mean (SD) | 1.35 (1.21) | 1.22 (0.87) | 1.38 (1.04) | 0.90 (0.74) | 0.9636 | 0.6961 | 0.0538 |
| **BBBi** | n=14 | n=8 | n=19 | n=23 |  |  |  |
| Mean (SD) | 0.005 (0.002) | 0.006 (0.003) | 0.006 (0.002) | 0.008 (0.004) | 0.7639 | 0.7975 | 0.1647 |
| **ATN group CSF data** |  |  |  |  |  |  |  |
|  | *Aβ42/40* | *p-Tau* | *t-Tau* |  |  |  |  |
| *Cutoff for positive values* | <0.139 | >65.5 pg/ml | >695.9 pg/ml |  |  |  |  |
| CSF AB 42/40 (A) mean (SD) | 0.19 (0.03) | 0.10 (0.02) | 0.10 (0.02) | 0.09 (0.02) |  |  |  |
| CSF pTau-181 (pg/ml) (T) mean (SD) | 37.1 (13.4) | 48.4 (9.5) | 113.1 (37.8) | 205.2 (123.0) |  |  |  |
| CSF tTau (pg/ml) (N) mean (SD) | 298.7 (136.0) | 324.3 (111.1) | 504.9 (114.5) | 1150.1 (400.5) |  |  |  |

**Supplementary Table II:** Outliers removed within disease groups per analyte. Outlier tests were performed using the ROUT method (Q=1%). Number of outliers removed per group and per analyte are shown.

| Immune factor | CN |  | MCI |  | AD |  |
| --- | --- | --- | --- | --- | --- | --- |
|  | # values | Outliers | # values | Outliers | # values | Outliers |
| sTREM2 | 88 | 7 | 37 | 3 | 75 | 5 |
| ***Immunoregulatory/ Pleiotropic*** |  |  |  |  |  |  |
| Fractalkine | 88 | 13 | 37 | 5 | 75 | 10 |
| MDC | 88 | 2 | 37 | 1 | 75 | 0 |
| IL-6 | 88 | 19 | 37 | 6 | 75 | 16 |
| IL-7 | 88 | 8 | 37 | 7 | 75 | 6 |
| IL-9 | 88 | 19 | 37 | 6 | 75 | 15 |
| TGF-α | 88 | 18 | 37 | 5 | 75 | 13 |
| ***Anti-inflammatory*** |  |  |  |  |  |  |
| FGF-2 | 88 | 11 | 37 | 5 | 75 | 10 |
| G-CSF | 88 | 6 | 37 | 5 | 75 | 7 |
| IL-1RA | 88 | 15 | 37 | 5 | 75 | 12 |
| IL-4 | 88 | 25 | 37 | 10 | 75 | 17 |
| IL-5 | 88 | 17 | 37 | 6 | 75 | 17 |
| IL-10 | 88 | 7 | 37 | 8 | 75 | 5 |
| IL-13 | 88 | 20 | 37 | 5 | 75 | 15 |
| ***Pro-inflammatory*** |  |  |  |  |  |  |
| EGF | 88 | 13 | 37 | 4 | 75 | 12 |
| Eotaxin | 88 | 2 | 37 | 0 | 75 | 4 |
| Flt-3L | 88 | 5 | 37 | 2 | 75 | 2 |
| GM-CSF | 88 | 8 | 37 | 5 | 75 | 11 |
| GRO | 88 | 4 | 37 | 1 | 75 | 9 |
| IFNα2 | 88 | 6 | 37 | 5 | 75 | 8 |
| IFNγ | 88 | 15 | 37 | 5 | 75 | 6 |
| IL-1α | 88 | 20 | 37 | 8 | 75 | 17 |
| IL-1β | 88 | 14 | 37 | 5 | 75 | 11 |
| IL-2 | 88 | 11 | 37 | 3 | 75 | 11 |
| IL-3 | 88 | 6 | 37 | 5 | 75 | 5 |
| IL-8 | 88 | 24 | 37 | 5 | 75 | 12 |
| IL-12P40 | 88 | 8 | 37 | 4 | 75 | 6 |
| IL-12P70 | 88 | 15 | 37 | 6 | 75 | 10 |
| IL-15 | 88 | 7 | 37 | 4 | 75 | 2 |
| IL-17A | 88 | 12 | 37 | 5 | 75 | 9 |
| IP-10 | 88 | 2 | 37 | 1 | 75 | 4 |
| MCP-1 | 88 | 1 | 37 | 3 | 75 | 0 |
| MCP-3 | 88 | 20 | 37 | 5 | 75 | 15 |
| MIP-1α | 88 | 8 | 37 | 3 | 75 | 5 |
| MIP-1β | 88 | 3 | 37 | 3 | 75 | 4 |
| sCD40L | 88 | 5 | 37 | 2 | 75 | 2 |
| TNFα | 88 | 2 | 37 | 0 | 75 | 2 |
| TNFβ | 88 | 24 | 37 | 7 | 75 | 17 |

**Supplementary Table III:** Outliers removed within ATN groups per analyte. Outlier tests were performed using the ROUT method (Q=1%). Number of outliers removed per group and per analyte are shown.

| Immune factor | A-T-N- |  | A+T-N- |  | A+T+N- |  | A+T+N+ |  |
| --- | --- | --- | --- | --- | --- | --- | --- | --- |
|  | # values | Outliers | # values | Outliers | # values | Outliers | # values | Outliers |
| STREM2 | 31 | 4 | 17 | 1 | 37 | 3 | 39 | 2 |
| ***Immunoregulatory/ Pleiotropic*** |  |  |  |  |  |  |  |  |
| Fractalkine | 31 | 3 | 17 | 0 | 37 | 6 | 39 | 3 |
| MDC | 31 | 1 | 17 | 0 | 37 | 0 | 39 | 0 |
| IL-6 | 31 | 6 | 17 | 4 | 37 | 8 | 39 | 8 |
| IL-7 | 31 | 5 | 17 | 1 | 37 | 7 | 39 | 4 |
| IL-9 | 31 | 5 | 17 | 4 | 37 | 8 | 39 | 7 |
| TGF-α | 31 | 4 | 17 | 1 | 37 | 5 | 39 | 7 |
| ***Anti-inflammatory*** |  |  |  |  |  |  |  |  |
| FGF-2 | 31 | 4 | 17 | 2 | 37 | 6 | 39 | 5 |
| G-CSF | 31 | 2 | 17 | 1 | 37 | 4 | 39 | 2 |
| IL-1RA | 31 | 3 | 17 | 3 | 37 | 5 | 39 | 4 |
| IL-4 | 31 | 7 | 17 | 5 | 37 | 9 | 39 | 10 |
| IL-5 | 31 | 4 | 17 | 3 | 37 | 5 | 39 | 8 |
| IL-10 | 31 | 4 | 17 | 1 | 37 | 3 | 39 | 4 |
| IL-13 | 31 | 4 | 17 | 3 | 37 | 6 | 39 | 6 |
| ***Pro-inflammatory*** |  |  |  |  |  |  |  |  |
| EGF | 31 | 4 | 17 | 3 | 37 | 6 | 39 | 5 |
| Eotaxin | 31 | 1 | 17 | 0 | 37 | 4 | 39 | 0 |
| Flt-3L | 31 | 2 | 17 | 0 | 37 | 2 | 39 | 2 |
| GM-CSF | 31 | 3 | 17 | 0 | 37 | 9 | 39 | 4 |
| GRO | 31 | 1 | 17 | 0 | 37 | 5 | 39 | 6 |
| IFNα2 | 31 | 2 | 17 | 1 | 37 | 6 | 39 | 2 |
| IFNγ | 31 | 6 | 17 | 3 | 37 | 4 | 39 | 6 |
| IL-1α | 31 | 6 | 17 | 4 | 37 | 8 | 39 | 8 |
| IL-1β | 31 | 4 | 17 | 2 | 37 | 4 | 39 | 5 |
| IL-2 | 31 | 3 | 17 | 2 | 37 | 3 | 39 | 7 |
| IL-3 | 31 | 4 | 17 | 0 | 37 | 4 | 39 | 1 |
| IL-8 | 31 | 4 | 17 | 2 | 37 | 7 | 39 | 6 |
| IL-12P40 | 31 | 3 | 17 | 0 | 37 | 2 | 39 | 3 |
| IL-12P70 | 31 | 6 | 17 | 4 | 37 | 5 | 39 | 5 |
| IL-15 | 31 | 1 | 17 | 0 | 37 | 4 | 39 | 2 |
| IL-17A | 31 | 4 | 17 | 3 | 37 | 5 | 39 | 4 |
| IP-10 | 31 | 0 | 17 | 0 | 37 | 1 | 39 | 3 |
| MCP-1 | 31 | 0 | 17 | 0 | 37 | 0 | 39 | 1 |
| MCP-3 | 31 | 7 | 17 | 2 | 37 | 7 | 39 | 11 |
| MIP-1α | 31 | 3 | 17 | 1 | 37 | 3 | 39 | 4 |
| MIP-1β | 31 | 0 | 17 | 2 | 37 | 3 | 39 | 0 |
| sCD40L | 31 | 1 | 17 | 0 | 37 | 1 | 39 | 1 |
| TNFα | 31 | 0 | 17 | 0 | 37 | 0 | 39 | 0 |
| TNFβ | 31 | 7 | 17 | 4 | 37 | 8 | 39 | 11 |
| VEGF | 31 | 4 | 17 | 4 | 37 | 4 | 39 | 5 |

**Supplementary Table IV:** Spearman correlations following outlier removal. Outlier removal was performed using the ROUT method (Q=1%). After outlier removal, sTREM2 and inflammatory factors were correlated within each disease and ATN group using the Spearman method. Spearman r correlation coefficients and p values are shown. Significant p values are bolded.

| Immune factor | CN |  | MCI |  | AD |  | A-T-N- |  | A+T-N- |  | A+T+N- |  | A+T+N+ |  |
| --- | --- | --- | --- | --- | --- | --- | --- | --- | --- | --- | --- | --- | --- | --- |
|  | Spearman r | p value | Spearman r | p value | Spearman r | p value | Spearman r | p value | Spearman r | p value | Spearman r | p value | Spearman r | p value |
| *Immunoregulatory/ Pleiotropic* |  |  |  |  |  |  |  |  |  |  |  |  |  |  |
| Fractalkine | 0.3035 | **0.0090** | 0.1451 | 0.4442 | 0.0382 | 0.7699 | 0.2944 | 0.1531 | 0.1844 | 0.5106 | 0.1466 | 0.4481 | 0.2314 | 0.1745 |
| MDC | 0.0466 | 0.6835 | 0.0128 | 0.9426 | 0.1470 | 0.2246 | 0.4188 | **0.0297** | 0.0054 | 0.9849 | 0.2483 | 0.1567 | -0.0377 | 0.8246 |
| IL-6 | 0.1603 | 0.1984 | -0.2154 | 0.2529 | 0.1679 | 0.2161 | 0.2519 | 0.2581 | -0.1210 | 0.6936 | 0.1820 | 0.3539 | 0.1828 | 0.3250 |
| IL-7 | 0.3479 | **0.0019** | -0.0374 | 0.8474 | 0.1872 | 0.1354 | 0.1027 | 0.6411 | -0.3819 | 0.1778 | 0.1778 | 0.3562 | 0.3565 | **0.0356** |
| IL-9 | 0.1861 | 0.1346 | -0.1715 | 0.3649 | -0.0366 | 0.7871 | 0.4080 | 0.0533 | -0.5144 | 0.0721 | 0.2661 | 0.1711 | 0.0016 | 0.9932 |
| TGF-α | 0.3046 | **0.0109** | 0.0333 | 0.8588 | 0.0745 | 0.5782 | 0.2186 | 0.3048 | -0.2599 | 0.3661 | 0.3692 | **0.0447** | 0.2846 | 0.1144 |
| *Anti-inflammatory* |  |  |  |  |  |  |  |  |  |  |  |  |  |  |
| FGF-2 | 0.3831 | **0.0008** | 0.1024 | 0.5902 | -0.0131 | 0.9202 | 0.1599 | 0.4553 | -0.2342 | 0.4376 | -0.0271 | 0.8889 | 0.3512 | **0.0417** |
| G-CSF | 0.2680 | **0.0177** | 0.0532 | 0.7799 | 0.2048 | 0.1045 | 0.0583 | 0.7818 | 0.2797 | 0.3302 | 0.2094 | 0.2583 | 0.2810 | 0.0969 |
| IL-1RA | -0.0296 | 0.8068 | 0.0249 | 0.8941 | 0.1009 | 0.4432 | -0.1120 | 0.5939 | -0.4251 | 0.1297 | 0.1777 | 0.3390 | 0.2152 | 0.2144 |
| IL-4 | 0.1763 | 0.1778 | -0.0017 | 0.9934 | -0.0843 | 0.5405 | 0.3955 | 0.0760 | -0.6737 | **0.0163** | 0.0634 | 0.7535 | 0.0581 | 0.7646 |
| IL-5 | 0.1895 | 0.1216 | -0.0686 | 0.7188 | -0.1619 | 0.2375 | 0.5469 | **0.0057** | -0.0397 | 0.8930 | 0.0989 | 0.5968 | -0.0318 | 0.8652 |
| IL-10 | 0.2469 | **0.0304** | 0.2781 | 0.1518 | 0.3233 | **0.0086** | 0.1506 | 0.4824 | 0.1036 | 0.7244 | 0.3924 | **0.0263** | 0.2534 | 0.1482 |
| IL-13 | -0.0047 | 0.9703 | -0.0591 | 0.7521 | 0.0305 | 0.8216 | 0.2416 | 0.2554 | -0.3476 | 0.2233 | 0.3445 | 0.0623 | -0.0041 | 0.9819 |
| *Pro-inflammatory* |  |  |  |  |  |  |  |  |  |  |  |  |  |  |
| EGF | 0.3202 | **0.0065** | 0.1297 | 0.4791 | 0.2760 | **0.0343** | 0.4629 | **0.0198** | 0.2273 | 0.4502 | 0.3903 | **0.0330** | 0.2888 | 0.1031 |
| Eotaxin | 0.1702 | 0.1312 | -0.0125 | 0.9439 | 0.0253 | 0.8403 | -0.3319 | 0.0977 | -0.1377 | 0.6222 | -0.0632 | 0.7401 | 0.0474 | 0.7804 |
| Flt-3L | 0.0923 | 0.4214 | -0.0342 | 0.8502 | 0.0826 | 0.5033 | 0.0466 | 0.8250 | -0.2587 | >0.9999 | 0.0831 | 0.6458 | -0.0179 | 0.9186 |
| GM-CSF | 0.3763 | **0.0007** | 0.1702 | 0.3599 | 0.1391 | 0.2890 | 0.3898 | 0.0541 | 0.0161 | 0.9545 | 0.1111 | 0.5813 | 0.3825 | **0.0233** |
| GRO | 0.1698 | 0.1347 | 0.1373 | 0.4462 | -0.0447 | 0.7304 | 0.2662 | 0.1887 | -0.0877 | 0.7561 | -0.1173 | 0.5446 | 0.4460 | **0.0105** |
| IFNα2 | 0.3916 | **0.0004** | -0.1558 | 0.4027 | 0.0595 | 0.6430 | 0.2339 | 0.2605 | -0.1652 | 0.5725 | -0.0549 | 0.7774 | 0.3576 | **0.0323** |
| IFNγ | 0.3362 | **0.0047** | -0.0820 | 0.6668 | 0.1587 | 0.2102 | 0.2545 | 0.2412 | 0.1261 | 0.6962 | -0.0877 | 0.6392 | 0.2641 | 0.1375 |
| IL-1α | 0.2095 | 0.0940 | 0.0500 | 0.7966 | 0.1151 | 0.4026 | 0.1739 | 0.4390 | 0.2393 | 0.4310 | 0.2346 | 0.2294 | 0.0257 | 0.8908 |
| IL-1β | 0.1764 | 0.1411 | 0.3186 | 0.0806 | 0.3437 | **0.0067** | 0.2311 | 0.2773 | -0.1996 | 0.4940 | 0.4537 | **0.0091** | 0.5759 | **0.0004** |
| IL-2 | 0.3609 | **0.0015** | 0.1075 | 0.5514 | 0.0101 | 0.9391 | 0.0358 | 0.8683 | 0.0331 | 0.9146 | -0.0031 | 0.9865 | 0.2611 | 0.1489 |
| IL-3 | 0.2142 | 0.0597 | 0.0004 | 0.9983 | 0.2471 | **0.0472** | -0.0682 | 0.7514 | -0.1782 | 0.5253 | 0.1567 | 0.3998 | 0.5442 | **0.0005** |
| IL-8 | 0.1833 | 0.1539 | -0.2608 | 0.1565 | 0.0430 | 0.7444 | 0.2962 | 0.1599 | -0.3184 | 0.2474 | 0.0764 | 0.6938 | -0.0050 | 0.9779 |
| IL-12P40 | 0.2084 | 0.0690 | 0.1011 | 0.5821 | 0.1354 | 0.2824 | 0.1182 | 0.5736 | -0.3160 | 0.2512 | 0.1783 | 0.3208 | 0.1860 | 0.2776 |
| IL-12P70 | 0.2119 | 0.0740 | 0.0890 | 0.6461 | 0.0553 | 0.6719 | 0.3120 | 0.1575 | -0.6164 | **0.0434** | -0.0770 | 0.6858 | 0.3095 | 0.0749 |
| IL-15 | 0.2159 | 0.0576 | -0.0209 | 0.9096 | 0.0655 | 0.5957 | 0.2750 | 0.1651 | -0.3987 | 0.1410 | 0.1797 | 0.3333 | 0.1630 | 0.3423 |
| IL-17A | 0.4163 | **0.0002** | -0.1111 | 0.5588 | 0.0943 | 0.4659 | 0.3359 | 0.1086 | 0.1331 | 0.6801 | 0.0613 | 0.7475 | 0.2758 | 0.1087 |
| IP-10 | 0.2951 | **0.0083** | -0.1932 | 0.2813 | 0.1995 | 0.1082 | 0.4801 | **0.0113** | 0.0555 | 0.8444 | 0.1473 | 0.4135 | 0.0841 | 0.6364 |
| MCP-1 | 0.1761 | 0.1181 | -0.0071 | 0.9699 | -0.0864 | 0.4767 | 0.0009 | 0.9964 | -0.1878 | 0.5026 | -0.3165 | 0.0682 | -0.0420 | 0.8080 |
| MCP-3 | 0.1956 | 0.1183 | -0.0689 | 0.7126 | 0.0652 | 0.6300 | 0.2783 | 0.2219 | -0.2880 | 0.2979 | 0.3192 | 0.0914 | 0.1656 | 0.3996 |
| MIP-1α | 0.2712 | **0.0170** | -0.0480 | 0.7908 | 0.2232 | 0.0716 | 0.0562 | 0.7895 | -0.0726 | 0.8052 | 0.1847 | 0.3114 | 0.0153 | 0.9303 |
| MIP-1β | -0.0686 | 0.5482 | 0.1918 | 0.2929 | 0.1722 | 0.1668 | 0.0254 | 0.9001 | -0.3912 | 0.1863 | 0.2416 | 0.1827 | 0.0314 | 0.8535 |
| sCD40L | 0.3451 | **0.0018** | -0.2771 | 0.1185 | 0.0180 | 0.8831 | 0.0926 | 0.6529 | -0.2113 | 0.4497 | -0.1816 | 0.3040 | 0.0637 | 0.7081 |
| TNFα | 0.3352 | **0.0022** | -0.2357 | 0.1796 | 0.1594 | 0.1943 | 0.2584 | 0.1931 | -0.0036 | 0.9899 | -0.1222 | 0.4911 | 0.2224 | 0.1859 |
| TNFβ | 0.1655 | 0.2024 | 0.0032 | 0.9868 | 0.0652 | 0.6363 | 0.1472 | 0.5243 | -0.3030 | 0.3142 | 0.2176 | 0.2661 | 0.3016 | 0.1188 |
| VEGF | 0.4491 | **<0.0001** | 0.0477 | 0.7991 | 0.3083 | **0.0156** | 0.4083 | 0.0531 | 0.3958 | 0.2028 | 0.3308 | 0.0691 | 0.3370 | 0.0551 |

**Supplementary Table V:** sTREM2 showed more significant relationships with inflammatory factors in CN than MCI and AD. Single linear regressions were run on sTREM2 with each cytokine within disease groups and ATN groups. Significant Beta coefficients are noted in bold.

| Immune factor | CN |  | MCI |  | AD |  | A-T-N- |  | A+T-N- |  | A+T+N- |  | A+T+N+ |  |
| --- | --- | --- | --- | --- | --- | --- | --- | --- | --- | --- | --- | --- | --- | --- |
|  | Beta coefficient | p value | Beta coefficient | p value | Beta coefficient | p value | Beta coefficient | p value | Beta coefficient | p value | Beta coefficient | p value | Beta coefficient | p value |
| **Immunoregulatory/ Pleiotropic** |  |  |  |  |  |  |  |  |  |  |  |  |  |  |
| Fractalkine | 0.2718 | **0.0304** | 0.2696 | **0.0489** | 0.0609 | 0.7093 | 0.2454 | 0.1321 | -0.2010 | 0.5310 | 0.1941 | 0.3868 | 0.3132 | 0.2974 |
| MDC | 0.0188 | 0.9972 | -0.0862 | 0.7825 | 0.1064 | 0.3899 | 0.0535 | 0.7365 | 0.6007 | **0.0389** | 0.1039 | 0.6291 | 0.1405 | 0.6320 |
| IL-6 | 0.1481 | 0.3810 | -0.1138 | 0.6393 | 0.1181 | 0.7175 | 0.1412 | 0.4257 | 0.1094 | 0.7635 | -0.1619 | 0.4952 | 0.1459 | 0.6509 |
| IL-7 | 0.3460 | **0.0085** | -0.0342 | 0.8518 | 0.1598 | 0.7694 | 0.1029 | 0.5445 | -0.2661 | 0.4888 | 0.0929 | 0.6811 | 0.5662 | **0.0348** |
| IL-9 | 0.1245 | 0.5775 | -0.1386 | 0.3036 | 0.0872 | 0.3758 | 0.2200 | 0.1906 | -0.3033 | 0.3943 | -0.0366 | 0.8818 | 0.3994 | 0.1984 |
| TGF-α | 0.3343 | **0.0066** | -0.1600 | 0.2913 | 0.0892 | 0.6091 | 0.2269 | 0.1769 | -0.2566 | 0.4462 | 0.0691 | 0.7721 | 0.2457 | 0.4414 |
| **Anti-inflammatory** |  |  |  |  |  |  |  |  |  |  |  |  |  |  |
| FGF-2 | 0.3472 | **0.1500** | 0.1569 | 0.4585 | -0.1315 | 0.2469 | 0.1314 | 0.4317 | -0.2837 | 0.3979 | 0.0597 | 0.7920 | 0.4027 | 0.1725 |
| G-CSF | 0.2939 | **0.0097** | 0.0849 | 0.5730 | 0.1046 | 0.9623 | 0.2257 | 0.1559 | -0.3031 | 0.3649 | -0.0368 | 0.8646 | 0.1081 | 0.7131 |
| IL-1RA | -0.0329 | 0.8900 | 0.0063 | 0.8640 | 0.1194 | 0.4642 | 0.0895 | 0.5828 | -0.3758 | 0.2845 | 0.0507 | 0.8227 | -0.1753 | 0.5489 |
| IL-4 | 0.1258 | 0.5472 | 0.0587 | 0.9635 | -0.0949 | 0.3731 | 0.1033 | 0.5673 | -0.2715 | 0.4797 | -0.2386 | 0.3252 | 0.1953 | 0.5430 |
| IL-5 | 0.0857 | 0.8667 | -0.0483 | 0.9177 | -0.1507 | 0.2449 | 0.4792 | **0.0027** | -0.2369 | 0.5099 | 0.0654 | 0.7783 | -0.0122 | 0.9684 |
| IL-10 | 0.3067 | **0.0033** | 0.1301 | 0.7320 | 0.2949 | **0.0124** | 0.2760 | 0.0934 | 0.7606 | **0.0106** | 0.4433 | **0.0441** | 0.0277 | 0.9251 |
| IL-13 | 0.0005 | 0.8757 | 0.0040 | 0.8766 | 0.0167 | 0.8567 | 0.2407 | 0.1513 | -0.1339 | 0.6947 | 0.1535 | 0.4952 | 0.0023 | 0.9949 |
| **Pro-inflammatory** |  |  |  |  |  |  |  |  |  |  |  |  |  |  |
| EGF | 0.3170 | **0.0134** | 0.0297 | 0.9279 | 0.3222 | **0.0056** | 0.3834 | **0.0146** | -0.0370 | 0.9192 | 0.2443 | 0.2612 | 0.5643 | **0.0445** |
| Eotaxin | 0.2044 | 0.1199 | 0.1107 | 0.5770 | 0.0917 | 0.5614 | -0.3043 | 0.0531 | 0.1226 | 0.7044 | 0.2012 | 0.3818 | 0.3552 | 0.2126 |
| Flt-3L | 0.0688 | 0.7924 | 0.1635 | 0.3876 | 0.0797 | 0.7523 | 0.0727 | 0.6515 | -0.2045 | 0.5238 | 0.2357 | 0.2790 | 0.3658 | 0.1983 |
| GM-CSF | 0.3839 | **0.0007** | 0.1069 | 0.6345 | 0.1652 | 0.1793 | 0.2683 | 0.1034 | -0.0222 | 0.9455 | -0.0139 | 0.9522 | 0.3210 | 0.2848 |
| GRO | 0.1675 | 0.1869 | 0.2550 | 0.5314 | -0.0291 | 0.5514 | 0.2348 | 0.1395 | 0.2856 | 0.3682 | -0.1922 | 0.4169 | 0.2546 | 0.4499 |
| IFNα2 | 0.4152 | **0.0003** | -0.0680 | 0.8029 | 0.1201 | 0.3360 | 0.1574 | 0.3453 | -0.3100 | 0.3267 | -0.0291 | 0.8953 | 0.2700 | 0.3505 |
| IFNγ | 0.3712 | **0.0010** | -0.0597 | 0.6916 | 0.0894 | 0.9747 | 0.2262 | 0.1722 | -0.3623 | 0.2735 | -0.2180 | 0.3424 | 0.4286 | 0.1440 |
| IL-1α | 0.2080 | 0.2013 | 0.0115 | 0.8681 | 0.1527 | 0.2965 | 0.1642 | 0.3458 | -0.2265 | 0.5579 | -0.1940 | 0.4124 | -0.0070 | 0.9827 |
| IL-1β | 0.1752 | 0.1889 | 0.0435 | 0.6625 | 0.4213 | **0.0005** | 0.2671 | 0.1002 | -0.2190 | 0.5713 | 0.3782 | 0.0827 | 0.1202 | 0.7099 |
| IL-2 | 0.3526 | **0.0026** | 0.0508 | 0.9644 | 0.0625 | 0.7615 | 0.2434 | 0.1408 | -0.4207 | 0.1976 | -0.1967 | 0.3684 | 0.2936 | 0.3808 |
| IL-3 | 0.1789 | 0.1915 | 0.0591 | 0.6229 | 0.3021 | **0.0132** | 0.1070 | 0.5112 | -0.2851 | 0.3692 | 0.1949 | 0.3728 | 0.2570 | 0.3752 |
| IL-8 | 0.1438 | 0.2777 | -0.0204 | 0.4199 | -0.0179 | 0.4237 | 0.2158 | 0.1996 | -0.2562 | 0.4470 | 0.0973 | 0.6748 | 0.1940 | 0.5255 |
| IL-12P40 | 0.2373 | 0.0870 | 0.1391 | 0.2526 | 0.0672 | 0.6538 | 0.1639 | 0.3058 | -0.2947 | 0.3525 | 0.0765 | 0.7350 | -0.1559 | 0.5947 |
| IL-12P70 | 0.2418 | 0.0618 | -0.0716 | 0.3214 | 0.0414 | 0.9717 | 0.0337 | 0.8453 | -0.2062 | 0.5676 | -0.0298 | 0.8928 | 0.4237 | 0.1941 |
| IL-15 | 0.2282 | 0.0848 | 0.0633 | 0.6399 | 0.0580 | 0.6149 | 0.1291 | 0.4211 | -0.2507 | 0.4318 | 0.2555 | 0.2637 | -0.0455 | 0.8773 |
| IL-17A | 0.4731 | **0.0000** | -0.1569 | 0.3590 | 0.1265 | 0.3698 | 0.3117 | 0.0642 | -0.5424 | 0.0848 | -0.1502 | 0.5047 | 0.4116 | 0.1437 |
| IP-10 | 0.2393 | 0.0805 | -0.2759 | 0.2022 | 0.1369 | 0.6427 | 0.3295 | **0.0354** | -0.2481 | 0.4619 | 0.1469 | 0.5142 | -0.1436 | 0.6242 |
| MCP-1 | 0.1268 | 0.4243 | -0.0206 | 0.8567 | 0.0135 | 0.3576 | 0.0094 | 0.9531 | 0.1149 | 0.7223 | -0.2433 | 0.2520 | -0.0011 | 0.9971 |
| MCP-3 | 0.1120 | 0.3167 | -0.0783 | 0.7345 | 0.0721 | 0.7378 | 0.4144 | **0.0148** | 0.0042 | 0.9902 | 0.1533 | 0.5071 | 0.2582 | 0.4177 |
| MIP-1α | 0.2821 | **0.0120** | -0.1452 | 0.3219 | 0.2076 | 0.0684 | 0.2346 | 0.1451 | -0.0641 | 0.8699 | 0.0278 | 0.8999 | -0.2667 | 0.3566 |
| MIP-1β | -0.0812 | 0.7751 | 0.2325 | 0.1136 | 0.1715 | 0.3320 | 0.2867 | 0.0768 | -0.2510 | 0.4843 | 0.3043 | 0.1483 | -0.3614 | 0.2042 |
| sCD40L | 0.2923 | **0.0062** | -0.1829 | 0.2718 | 0.0199 | 0.9306 | -0.0763 | 0.6398 | -0.1773 | 0.5815 | -0.1009 | 0.6390 | -0.0981 | 0.7387 |
| TNFα | 0.2730 | **0.0251** | -0.1566 | 0.3967 | 0.2183 | 0.2132 | 0.3895 | **0.0108** | -0.2808 | 0.3767 | -0.2024 | 0.3428 | -0.0428 | 0.8846 |
| TNFβ | 0.1358 | 0.4718 | -0.0785 | 0.7768 | -0.0040 | 0.4986 | -0.0146 | 0.9366 | -0.1111 | 0.7759 | -0.1342 | 0.5727 | 0.1056 | 0.7439 |
| VEGF | 0.4457 | **0.0002** | 0.1229 | 0.3711 | 0.2552 | 0.0721 | 0.2124 | 0.2068 | 0.1204 | 0.7403 | 0.1760 | 0.4580 | 0.3813 | 0.2214 |

**Supplementary Table VI:** Outliers removed within sTREM2 Tertiles 1 and 3, per analyte. Outlier tests were performed using the ROUT method (Q=1%). Number of outliers removed per group and per analyte are shown.

| Immune factor | sTREM2 Tertile 1 |  | sTREM2 Tertile 3 |  |
| --- | --- | --- | --- | --- |
|  | # values | Outliers | # values | Outliers |
| sTREM2 | 67 | 0 | 67 | 6 |
| ***Immunoregulatory/ Pleiotropic*** |  |  |  |  |
| Fractalkine | 67 | 8 | 67 | 10 |
| MDC | 67 | 1 | 67 | 1 |
| IL-6 | 67 | 12 | 67 | 14 |
| IL-7 | 67 | 6 | 67 | 8 |
| IL-9 | 67 | 15 | 67 | 18 |
| TGF-α | 67 | 6 | 67 | 15 |
| ***Anti-inflammatory*** |  |  |  |  |
| FGF-2 | 67 | 7 | 67 | 9 |
| G-CSF | 67 | 7 | 67 | 6 |
| IL-1RA | 67 | 4 | 67 | 15 |
| IL-4 | 67 | 17 | 67 | 10 |
| IL-5 | 67 | 13 | 67 | 14 |
| IL-10 | 67 | 4 | 67 | 8 |
| IL-13 | 67 | 11 | 67 | 15 |
| ***Pro-inflammatory*** |  |  |  |  |
| EGF | 67 | 5 | 67 | 8 |
| Eotaxin | 67 | 3 | 67 | 3 |
| Flt-3L | 67 | 4 | 67 | 9 |
| GM-CSF | 67 | 5 | 67 | 5 |
| GRO | 67 | 5 | 67 | 1 |
| IFNα2 | 67 | 4 | 67 | 5 |
| IFNγ | 67 | 10 | 67 | 11 |
| IL-1α | 67 | 15 | 67 | 8 |
| IL-1β | 67 | 11 | 67 | 6 |
| IL-2 | 67 | 8 | 67 | 9 |
| IL-3 | 67 | 4 | 67 | 3 |
| IL-8 | 67 | 10 | 67 | 8 |
| IL-12P40 | 67 | 4 | 67 | 12 |
| IL-12P70 | 67 | 12 | 67 | 13 |
| IL-15 | 67 | 2 | 67 | 8 |
| IL-17A | 67 | 10 | 67 | 10 |
| IP-10 | 67 | 2 | 67 | 3 |
| MCP-1 | 67 | 1 | 67 | 2 |
| MCP-3 | 67 | 11 | 67 | 14 |
| MIP-1α | 67 | 4 | 67 | 8 |
| MIP-1β | 67 | 4 | 67 | 3 |
| sCD40L | 67 | 3 | 67 | 5 |
| TNFα | 67 | 2 | 67 | 2 |
| TNFβ | 67 | 13 | 67 | 10 |

**Supplementary Table VII:**
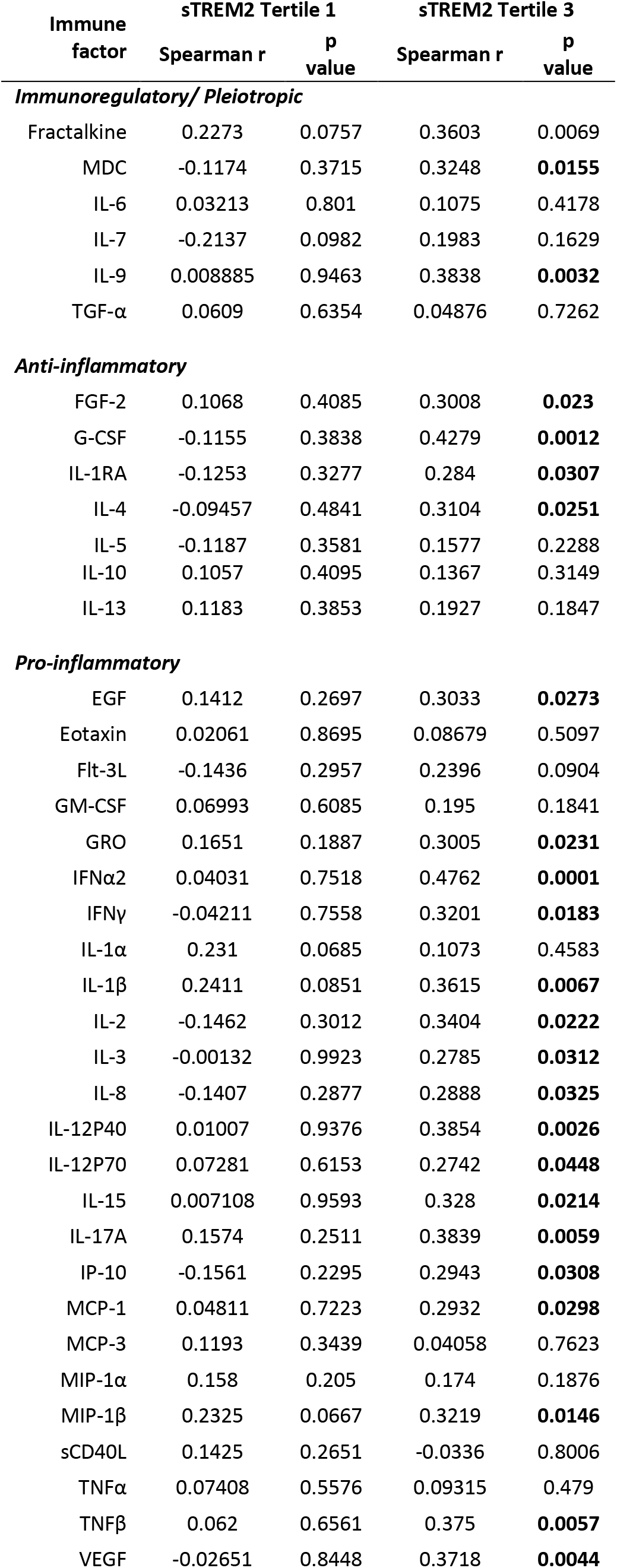
Spearman correlations following outlier removal within sTREM2 Tertiles 1 and 3. Outlier removal was performed using the ROUT method (Q=1%). After outlier removal, sTREM2 and inflammatory factors were correlated within sTREM2 tertiles 1 and 3 using the Spearman method. Spearman r correlation coefficients and p values are shown. Significant p values are bolded

## References

1. 2020 Alzheimer’s disease facts and figures. Alzheimers Dement. 2020.

2. Zetterberg H. Blood-based biomarkers for Alzheimer’s disease-An update. J Neurosci Methods. 2019;319:2–6.

3. Molinuevo J. L., Ayton S., Batrla R., Bednar M. M., Bittner T., Cummings J., Fagan A. M., Hampel H., Mielke M. M., Mikulskis A., O’Bryant S., Scheltens P., Sevigny J., Shaw L. M., Soares H. D., Tong G., Trojanowski J. Q., Zetterberg H., Blennow K. Current state of Alzheimer’s fluid biomarkers. Acta Neuropathol. 2018;136(6):821–53.

4. Dubois B., Feldman H. H., Jacova C., Hampel H., Molinuevo J. L., Blennow K., DeKosky S. T., Gauthier S., Selkoe D., Bateman R., Cappa S., Crutch S., Engelborghs S., Frisoni G. B., Fox N. C., Galasko D., Habert M. O., Jicha G. A., Nordberg A., Pasquier F., Rabinovici G., Robert P., Rowe C., Salloway S., Sarazin M., Epelbaum S., de Souza L. C., Vellas B., Visser P. J., Schneider L., Stern Y., Scheltens P., Cummings J. L. Advancing research diagnostic criteria for Alzheimer’s disease: the IWG-2 criteria. Lancet Neurol. 2014;13(6):614–29.

5. Zetterberg H. Applying fluid biomarkers to Alzheimer’s disease. Am J Physiol Cell Physiol. 2017;313(1):C3–C10.

6. Kidd P. M. Alzheimer’s disease, amnestic mild cognitive impairment, and age-associated memory impairment: current understanding and progress toward integrative prevention. Altern Med Rev. 2008;13(2):85–115.

7. Davis M., T O. C., Johnson S., Cline S., Merikle E., Martenyi F., Simpson K. Estimating Alzheimer’s Disease Progression Rates from Normal Cognition Through Mild Cognitive Impairment and Stages of Dementia. Curr Alzheimer Res. 2018;15(8):777–88.

8. Jack C. R., Jr., Bennett D. A., Blennow K., Carrillo M. C., Feldman H. H., Frisoni G. B., Hampel H., Jagust W. J., Johnson K. A., Knopman D. S., Petersen R. C., Scheltens P., Sperling R. A., Dubois B. A/T/N: An unbiased descriptive classification scheme for Alzheimer disease biomarkers. Neurology. 2016;87(5):539–47.

9. Calsolaro V., Edison P. Neuroinflammation in Alzheimer’s disease: Current evidence and future directions. Alzheimers Dement. 2016;12(6):719–32.

10. Le Page A., Dupuis G., Frost E. H., Larbi A., Pawelec G., Witkowski J. M., Fulop T. Role of the peripheral innate immune system in the development of Alzheimer’s disease. Exp Gerontol. 2018;107:59–66.

11. Wotton C. J., Goldacre M. J. Associations between specific autoimmune diseases and subsequent dementia: retrospective record-linkage cohort study, UK. J Epidemiol Community Health. 2017;71(6):576–83.

12. Costantini E., D’Angelo C., Reale M. The Role of Immunosenescence in Neurodegenerative Diseases. Mediators Inflamm. 2018;2018:6039171.

13. Ng A., Tam W. W., Zhang M. W., Ho C. S., Husain S. F., McIntyre R. S., Ho R. C. IL-1beta, IL-6, TNF-alpha and CRP in Elderly Patients with Depression or Alzheimer’s disease: Systematic Review and Meta-Analysis. Sci Rep. 2018;8(1):12050.

14. Su C., Zhao K., Xia H., Xu Y. Peripheral inflammatory biomarkers in Alzheimer’s disease and mild cognitive impairment: a systematic review and meta-analysis. Psychogeriatrics. 2019;19(4):300–9.

15. Swardfager W., Lanctot K., Rothenburg L., Wong A., Cappell J., Herrmann N. A meta-analysis of cytokines in Alzheimer’s disease. Biol Psychiatry. 2010;68(10):930–41.

16. Guerreiro R., Wojtas A., Bras J., Carrasquillo M., Rogaeva E., Majounie E., Cruchaga C., Sassi C., Kauwe J. S., Younkin S., Hazrati L., Collinge J., Pocock J., Lashley T., Williams J., Lambert J. C., Amouyel P., Goate A., Rademakers R., Morgan K., Powell J., St George-Hyslop P., Singleton A., Hardy J., Alzheimer Genetic Analysis G. TREM2 variants in Alzheimer’s disease. N Engl J Med. 2013;368(2):117–27.

17. Jonsson T., Stefansson H., Steinberg S., Jonsdottir I., Jonsson P. V., Snaedal J., Bjornsson S., Huttenlocher J., Levey A. I., Lah J. J., Rujescu D., Hampel H., Giegling I., Andreassen O. A., Engedal K., Ulstein I., Djurovic S., Ibrahim-Verbaas C., Hofman A., Ikram M. A., van Duijn C. M., Thorsteinsdottir U., Kong A., Stefansson K. Variant of TREM2 associated with the risk of Alzheimer’s disease. N Engl J Med. 2013;368(2):107–16.

18. Schlepckow K., Kleinberger G., Fukumori A., Feederle R., Lichtenthaler S. F., Steiner H., Haass C. An Alzheimer-associated TREM2 variant occurs at the ADAM cleavage site and affects shedding and phagocytic function. EMBO Mol Med. 2017;9(10):1356–65.

19. Thornton P., Sevalle J., Deery M. J., Fraser G., Zhou Y., Stahl S., Franssen E. H., Dodd R. B., Qamar S., Gomez Perez-Nievas B., Nicol L. S., Eketjall S., Revell J., Jones C., Billinton A., St George-Hyslop P. H., Chessell I., Crowther D. C. TREM2 shedding by cleavage at the H157-S158 bond is accelerated for the Alzheimer’s disease-associated H157Y variant. EMBO Mol Med. 2017;9(10):1366–78.

20. Liu D., Cao B., Zhao Y., Huang H., McIntyre R. S., Rosenblat J. D., Zhou H. Soluble TREM2 changes during the clinical course of Alzheimer’s disease: A meta-analysis. Neurosci Lett. 2018;686:10–6.

21. Shen X. N., Niu L. D., Wang Y. J., Cao X. P., Liu Q., Tan L., Zhang C., Yu J. T. Inflammatory markers in Alzheimer’s disease and mild cognitive impairment: a meta-analysis and systematic review of 170 studies. J Neurol Neurosurg Psychiatry. 2019;90(5):590–8.

22. Peskind E. R., Leverenz J., Farlow M. R., Ito R. K., Provow S. A., Siegel R. S., Cleveland M., Morgan C. H., Pandian M. R., Corbin S., Nochlin D., Schellenberg G. D., Raskind M. A., Wagner S. L. Clinicopathologic correlations of soluble amyloid beta-protein precursor in cerebrospinal fluid in patients with Alzheimer disease and controls. Alzheimer Dis Assoc Disord. 1997;11(4):201–6.

23. Bekris L. M., Millard S. P., Galloway N. M., Vuletic S., Albers J. J., Li G., Galasko D. R., DeCarli C., Farlow M. R., Clark C. M., Quinn J. F., Kaye J. A., Schellenberg G. D., Tsuang D., Peskind E. R., Yu C. E. Multiple SNPs within and surrounding the apolipoprotein E gene influence cerebrospinal fluid apolipoprotein E protein levels. J Alzheimers Dis. 2008;13(3):255–66.

24. Bekris L. M., Khrestian M., Dyne E., Shao Y., Pillai J. A., Rao S. M., Bemiller S. M., Lamb B., Fernandez H. H., Leverenz J. B. Soluble TREM2 and biomarkers of central and peripheral inflammation in neurodegenerative disease. J Neuroimmunol. 2018;319:19–27.

25. Breen E. J., Polaskova V., Khan A. Bead-based multiplex immuno-assays for cytokines, chemokines, growth factors and other analytes: median fluorescence intensities versus their derived absolute concentration values for statistical analysis. Cytokine. 2015;71(2):188–98.

26. Breen E. J., Tan W., Khan A. The Statistical Value of Raw Fluorescence Signal in Luminex xMAP Based Multiplex Immunoassays. Sci Rep. 2016;6:26996.

27. Helsel D. R. Fabricating data: how substituting values for nondetects can ruin results, and what can be done about it. Chemosphere. 2006;65(11):2434–9.

28. Kern S., Zetterberg H., Kern J., Zettergren A., Waern M., Hoglund K., Andreasson U., Wetterberg H., Borjesson-Hanson A., Blennow K., Skoog I. Prevalence of preclinical Alzheimer disease: Comparison of current classification systems. Neurology. 2018;90(19):e1682–e91.

29. Hoglund K., Kern S., Zettergren A., Borjesson-Hansson A., Zetterberg H., Skoog I., Blennow K. Preclinical amyloid pathology biomarker positivity: effects on tau pathology and neurodegeneration. Transl Psychiatry. 2017;7(1):e995.

30. Grontvedt G. R., Lauridsen C., Berge G., White L. R., Salvesen O., Brathen G., Sando S. B. The Amyloid, Tau, and Neurodegeneration (A/T/N) Classification Applied to a Clinical Research Cohort with Long-Term Follow-Up. J Alzheimers Dis. 2020;74(3):829–37.

31. Ben-Shachar M L. D., Makowski D. Estimation of Effect Size Indices and Standardized Parameters. Journal of Open Source Software. 2020;5(56):2815.

32. Taiyun Wei V. S. R package “corrplot”: Visualization of a Correlation Matrix Github 2017 [Version 0.84:[Available from: https://github.com/taiyun/corrplot.

33. Jr F. E. H. Hmisc: Harrell Miscellaneous 2021-02-28 [Available from: https://hbiostat.org/R/Hmisc/, https://github.com/harrelfe/Hmisc/.

34. Revelle W. psych: Procedures for Psychological, Psychometric, and Personality Research Evanston, Illinois: Northwestern University; 2020 [R package version 2.0.12:[Available from: https://cran.r-project.org/package=psych.

35. Bastian M., Heymann S., Jacomy M. Gephi: An Open Source Software for Exploring and Manipulating Networks. International AAAI Conference on Weblogs and Social Media 2009.

36. Ewers M., Franzmeier N., Suarez-Calvet M., Morenas-Rodriguez E., Caballero M. A. A., Kleinberger G., Piccio L., Cruchaga C., Deming Y., Dichgans M., Trojanowski J. Q., Shaw L. M., Weiner M. W., Haass C., Alzheimer’s Disease Neuroimaging I. Increased soluble TREM2 in cerebrospinal fluid is associated with reduced cognitive and clinical decline in Alzheimer’s disease. Sci Transl Med. 2019;11(507).

37. Leyns C. E. G., Gratuze M., Narasimhan S., Jain N., Koscal L. J., Jiang H., Manis M., Colonna M., Lee V. M. Y., Ulrich J. D., Holtzman D. M. TREM2 function impedes tau seeding in neuritic plaques. Nat Neurosci. 2019;22(8):1217–22.

38. Lee S. H., Meilandt W. J., Xie L., Gandham V. D., Ngu H., Barck K. H., Rezzonico M. G., Imperio J., Lalehzadeh G., Huntley M. A., Stark K. L., Foreman O., Carano R. A. D., Friedman B. A., Sheng M., Easton A., Bohlen C. J., Hansen D. V. Trem2 restrains the enhancement of tau accumulation and neurodegeneration by beta-amyloid pathology. Neuron. 2021;109(8):1283–301 e6.

39. Morgan A. R., Touchard S., Leckey C., O’Hagan C., Nevado-Holgado A. J., Consortium N., Barkhof F., Bertram L., Blin O., Bos I., Dobricic V., Engelborghs S., Frisoni G., Frolich L., Gabel S., Johannsen P., Kettunen P., Kloszewska I., Legido-Quigley C., Lleo A., Martinez-Lage P., Mecocci P., Meersmans K., Molinuevo J. L., Peyratout G., Popp J., Richardson J., Sala I., Scheltens P., Streffer J., Soininen H., Tainta-Cuezva M., Teunissen C., Tsolaki M., Vandenberghe R., Visser P. J., Vos S., Wahlund L. O., Wallin A., Westwood S., Zetterberg H., Lovestone S., Morgan B. P., Annex N.-W. T. C. f. N. o. M. D., Alzheimer’s D. Inflammatory biomarkers in Alzheimer’s disease plasma. Alzheimers Dement. 2019;15(6):776–87.

40. Azizi G., Khannazer N., Mirshafiey A. The Potential Role of Chemokines in Alzheimer’s Disease Pathogenesis. Am J Alzheimers Dis Other Demen. 2014;29(5):415–25.

41. Finneran D. J., Nash K. R. Neuroinflammation and fractalkine signaling in Alzheimer’s disease. J Neuroinflammation. 2019;16(1):30.

42. Zujovic V., Benavides J., Vige X., Carter C., Taupin V. Fractalkine modulates TNF-alpha secretion and neurotoxicity induced by microglial activation. Glia. 2000;29(4):305–15.

43. Desforges N. M., Hebron M. L., Algarzae N. K., Lonskaya I., Moussa C. E. Fractalkine Mediates Communication between Pathogenic Proteins and Microglia: Implications of Anti-Inflammatory Treatments in Different Stages of Neurodegenerative Diseases. Int J Alzheimers Dis. 2012;2012:345472.

44. Chen P., Zhao W., Guo Y., Xu J., Yin M. CX3CL1/CX3CR1 in Alzheimer’s Disease: A Target for Neuroprotection. Biomed Res Int. 2016;2016:8090918.

45. Pawelec P., Ziemka-Nalecz M., Sypecka J., Zalewska T. The Impact of the CX3CL1/CX3CR1 Axis in Neurological Disorders. Cells. 2020;9(10).

46. Strobel S., Grunblatt E., Riederer P., Heinsen H., Arzberger T., Al-Sarraj S., Troakes C., Ferrer I., Monoranu C. M. Changes in the expression of genes related to neuroinflammation over the course of sporadic Alzheimer’s disease progression: CX3CL1, TREM2, and PPARgamma. J Neural Transm (Vienna). 2015;122(7):1069–76.

47. Hundhausen C., Misztela D., Berkhout T. A., Broadway N., Saftig P., Reiss K., Hartmann D., Fahrenholz F., Postina R., Matthews V., Kallen K. J., Rose-John S., Ludwig A. The disintegrin-like metalloproteinase ADAM10 is involved in constitutive cleavage of CX3CL1 (fractalkine) and regulates CX3CL1-mediated cell-cell adhesion. Blood. 2003;102(4):1186–95.

48. Wood L. B., Winslow A. R., Proctor E. A., McGuone D., Mordes D. A., Frosch M. P., Hyman B. T., Lauffenburger D. A., Haigis K. M. Identification of neurotoxic cytokines by profiling Alzheimer’s disease tissues and neuron culture viability screening. Sci Rep. 2015;5:16622.

49. Zhou Y., Li C., Li D., Zheng Y., Wang J. IL-5 blocks apoptosis and tau hyperphosphorylation induced by Abeta25-35 peptide in PC12 cells. J Physiol Biochem. 2017;73(2):259–66.

50. Richartz E., Stransky E., Batra A., Simon P., Lewczuk P., Buchkremer G., Bartels M., Schott K. Decline of immune responsiveness: a pathogenetic factor in Alzheimer’s disease? J Psychiatr Res. 2005;39(5):535–43.

51. Italiani P., Puxeddu I., Napoletano S., Scala E., Melillo D., Manocchio S., Angiolillo A., Migliorini P., Boraschi D., Vitale E., Di Costanzo A. Circulating levels of IL-1 family cytokines and receptors in Alzheimer’s disease: new markers of disease progression? J Neuroinflammation. 2018;15(1):342.

52. Shaftel S. S., Kyrkanides S., Olschowka J. A., Miller J. N., Johnson R. E., O’Banion M. K. Sustained hippocampal IL-1 beta overexpression mediates chronic neuroinflammation and ameliorates Alzheimer plaque pathology. J Clin Invest. 2007;117(6):1595–604.

53. Lee B., Kim T. H., Jun J. B., Yoo D. H., Woo J. H., Choi S. J., Lee Y. H., Song G. G., Sohn J., Park-Min K. H., Ivashkiv L. B., Ji J. D. Direct inhibition of human RANK+ osteoclast precursors identifies a homeostatic function of IL-1beta. J Immunol. 2010;185(10):5926–34.

54. Gu C., Wu L., Li X. IL-17 family: cytokines, receptors and signaling. Cytokine. 2013;64(2):477–85.

55. Solleiro-Villavicencio H., Hernandez-Orozco E., Rivas-Arancibia S. Effect of exposure to low doses of ozone on interleukin 17A expression during progressive neurodegeneration in the rat hippocampus. Neurologia. 2018.

56. Cipollini V., Anrather J., Orzi F., Iadecola C. Th17 and Cognitive Impairment: Possible Mechanisms of Action. Front Neuroanat. 2019;13:95.

57. Delmotte K., Schaeverbeke J., Poesen K., Vandenberghe R. Prognostic value of amyloid/tau/neurodegeneration (ATN) classification based on diagnostic cerebrospinal fluid samples for Alzheimer’s disease. Alzheimers Res Ther. 2021;13(1):84.

58. Mila-Aloma M., Salvado G., Gispert J. D., Vilor-Tejedor N., Grau-Rivera O., Sala-Vila A., Sanchez-Benavides G., Arenaza-Urquijo E. M., Crous-Bou M., Gonzalez-de-Echavarri J. M., Minguillon C., Fauria K., Simon M., Kollmorgen G., Zetterberg H., Blennow K., Suarez-Calvet M., Molinuevo J. L., study A. Amyloid beta, tau, synaptic, neurodegeneration, and glial biomarkers in the preclinical stage of the Alzheimer’s continuum. Alzheimers Dement. 2020;16(10):1358–71.

59. Cousins K. A. Q., Phillips J. S., Irwin D. J., Lee E. B., Wolk D. A., Shaw L. M., Zetterberg H., Blennow K., Burke S. E., Kinney N. G., Gibbons G. S., McMillan C. T., Trojanowski J. Q., Grossman M. ATN incorporating cerebrospinal fluid neurofilament light chain detects frontotemporal lobar degeneration. Alzheimers Dement. 2021;17(5):822–30.

